# CRISPR-based targeted haplotype-resolved assemblies of a megabase region

**DOI:** 10.1101/2022.01.21.477044

**Authors:** Taotao Li, Duo Du, Dandan Zhang, Jiakang Ma, Mengyu Zhou, Weida Meng, Zelin Jin, Yicheng Lin, Ziqiang Chen, Haozhe Yuan, Jue Wang, Shulong Dong, Shaoyang Sun, Wenjing Ye, Boshen Li, Zhao Zhang, Zhi Xie, Wenqing Qiu, Yun Liu

## Abstract

Constructing high-quality haplotype-resolved genome assemblies has substantially improved the ability to detect and characterize genetic variants. A targeted approach providing readily access to the rich information from haplotype-resolved genome assemblies will be appealing to groups of basic researchers and medical scientists focused on specific genomic regions. Here, using the 4.5 megabase, notoriously difficult-to-assemble major histocompatibility complex (MHC) region as an example, we demonstrated an approach to construct haplotype-resolved *de novo* assemblies of targeted genomic regions with the CRISPR-based enrichment. Compared to the results from haplotype-resolved genome assemblies, our targeted approach achieved comparable completeness and accuracy with greatly reduced computing complexity, sequencing cost, as well as the amount of starting materials. Moreover, using the targeted assembled personal haplotypes as the reference both improves the quantification accuracy for sequencing data and enables allele-specific functional genomics analyses. Given its highly efficient use of resources, our approach can greatly facilitate population genetic studies of targeted regions, and may pave a new way to elucidate the molecular mechanisms in disease etiology.

---

The recent advances of constructing high-quality haplotype-resolved genome assemblies have substantially improved sensitivity in detecting and characterizing genetic variants^1^, and will greatly advance our understanding on the human genome. When parent-child trio information is available, the trio-binning approach permits the construction of haplotype-resolved genome assemblies for offspring individuals^2-4^. Alternatively, the PacBio high-fidelity (HiFi) long-read sequencing, coupled with strand-specific sequencing technologies, provides another option to assemble the diploid genome without the need for pedigree information^1, 5-8^. However, both approaches to achieve high-quality haplotype-resolved *de novo* genome assemblies require substantial large amounts of starting materials, extensive computing resources and high level of sequencing costs. A targeted approach providing readily access to the rich information from haplotype-resolved genome assemblies will be appealing to groups of basic researchers and medical scientists, who are focused on specific genomic regions for their functional relevance.

Recently, CRISPR/Cas9 starts to be adopted as a tool for the enrichment of targeted genomic regions^9-13^. Compared with probe-based DNA enrichment strategies, this approach only needs to know the DNA sequences flanking the targeted genomic region, making it an ideal tool for unbiased enrichment of targeted regions, even with high polymorphisms. Moreover, it is compatible with long-read sequencing technologies^9-13^ required for haplotype-resolved assemblies. We envisioned that adopting this targeted approach of the CRISPR-based enrichment could greatly reduce the required input and resources for achieving haplotype-resolved assemblies. To demonstrate the possibility, we chose the 4.5 megabase (Mb), notoriously difficult-to-assemble Major Histocompatibility Complex (MHC) region, which contains many genes directly involved in immune responses to antigens and is associated with many complex human diseases^14-16^. The studies of the MHC region are hampered by its extremely high polymorphisms and strong linkage disequilibrium. Achieving haplotype-resolved *de novo* assemblies of the MHC region is vital for unbiased detection and characterization of genetic variants in the MHC region^17^ for accurate genome inference^18^ towards the eventual understanding of its molecular mechanisms in disease etiology.

In this work, by combining the CRISPR-based enrichment with 10x Genomics linked-read sequencing and the PacBio HiFi long-read platform (Fig. 1a), we were able to achieve targeted haplotype-resolved *de novo* assemblies of the MHC region from diploid human cells. By comparing to the haplotype-resolved genome assemblies from the same cell line, we showed that our targeted approach for obtaining haplotype-resolved assemblies achieved comparable completeness and accuracy, enabling comprehensive detection of genetic variants of the targeted region. With the targeted assembled personal MHC haplotypes available as the reference, we showed that the quantifications of DNA methylation and gene expression of the MHC region are much more accurate, compared to the standard approach using the hg38 as the reference. Finally, we used the targeted assembled personal MHC haplotypes to investigate allele-specific transcriptional regulation of the MHC region.

**Figure 1.**
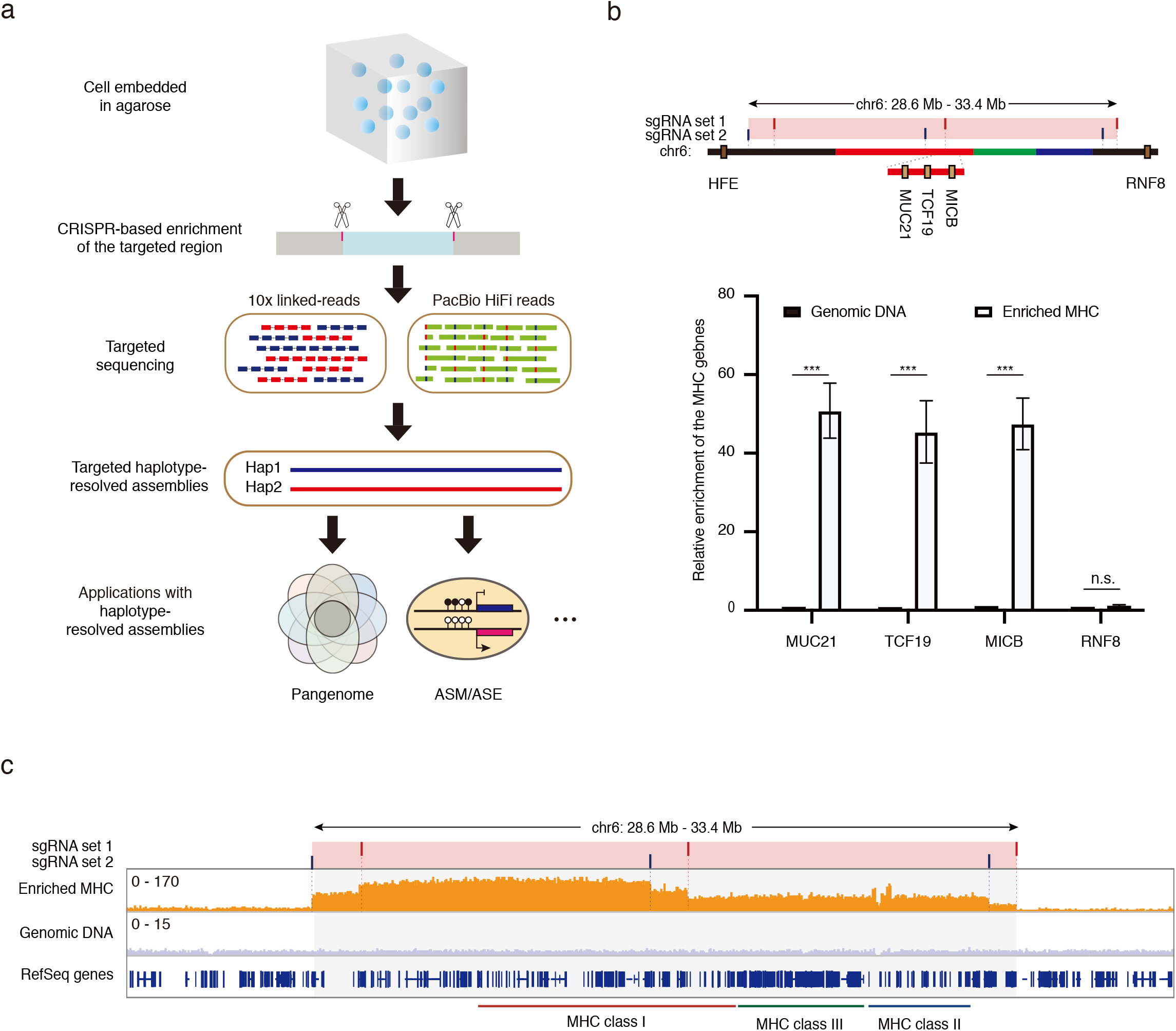
CRISPR-based targeted enrichment of the MHC region. **a**. Schematics of CRISPR-based targeted haplotype-resolved assemblies of a megabase region. Cells were first prepared and embedded in agarose plugs. The targeted genomic region was then enriched from agarose plugs by combining CRISPR-based in-gel digestion with pulsed field gel electrophoresis. Enriched HMW DNA molecules were subjected to both 10x Genomics linked-read and the PacBio HiFi long-read sequencing. By the intersection of variants generated from 10x Genomics linked-read data and HiFi reads, a highly reliable set of phased heterozygous variants was obtained to separate the HiFi reads into two haplotype-partitioned read sets, from which haplotype-resolved assemblies of the targeted region were constructed. With the targeted assembled personal haplotypes available, it can be widely used in many situations, including generating the haplotype-resolved pangenome, analyzing allele-specific methylation (ASM) or/and allele-specific expression (ASE), *et al*. **b**. QPCR analyses showed significant enrichment of the targeted MHC region. Top part: the positions of two sets of sgRNAs targeting the MHC region. The pink bar indicates the targeted MHC region. The red, green and blue lines indicate the regions for the HLA class I, class III and class II genes, respectively. The dark yellow boxes represent genes used for qPCR analyses. Genes are listed in order based on their coordinates on the hg38 reference but not to scale. Bottom part: the relative enrichment was determined relative to that from the *HFE* gene (indicated in the top part) outside of the targeted MHC region, and then normalized to that from cells treated with no sgRNA. The relative enrichment for another gene outside of the targeted MHC region (*RNF8*) was also tested as the negative control. Data are represented as mean ± SEM from three independent experiments (\*\*\**P* value < 0.001, student’s t-test, two-sided). **c**. The coverage of the targeted MHC region (the grey area) based on sequencing with the Illumina short-read platform. The targeted sites for two sets of sgRNAs are indicated as red and blue bars, respectively.

## Results

### CRISPR-based targeted enrichment of the MHC region

Our studies employed the well-characterized and widely used human diploid cell line, GM12878. The first step of our approach is the isolation of the targeted megabase-size DNA region (the MHC region) from agarose-embedded, intact cells by combining CRISPR-based in-gel digestion with pulsed field gel electrophoresis (Supplementary Fig.1a). SgRNAs were designed to target the non-polymorphic DNA sequences flanking the 4.7 Mb region known to include the entire MHC locus^19^ (Fig.1b). The performance of the designed sgRNAs was initially evaluated through *in vitro* cleavage assays with PCR products amplified from the targeted regions (Supplementary Fig. 1b). Most of the final enriched DNA molecules were still high molecular weight (HMW) with length longer than 50 kb (Supplementary Fig. 2), and are thus suitable for subsequent long-read sequencing.

To evaluate the enrichment efficiency, we performed qPCR analyses of three genomic loci within the targeted MHC region (chr6: 28903952-33268517), which revealed more than 40-fold enrichment compared to the no sgRNA controls; note that no enrichment was detected for the MHC-flanking region (Fig. 1b). Sequencing with the Illumina short-read platform showed that the whole targeted MHC region was successfully enriched (Fig. 1c). These results showed that, through the CRISPR/Cas9 cleavage followed by gel purification, we can successfully enrich the targeted MHC region.

### Targeted haplotype-resolved assemblies of the MHC region

After the enrichment of the MHC region, we constructed 10x Genomics linked-read libraries with the HMW MHC molecules. The phased variants within the targeted MHC region were called (Supplementary Fig. 3), and then compared to two benchmark datasets for GM12878 cells (the GIAB (v3.3.2)^20^ and the Illumina Platinum Genomes^21^). Most of the variants were identified with false negative rates (FNRs) lower than 4% (Table 1). The accuracy of phasing was high, with both switch error rates and Hamming error rates lower than 2% (Supplementary Table 1). We estimated the HLA types with HLA-VBSeq^22^ (Supplementary Fig. 3) for six major *HLA* genes, which have been found to be strongly associated with many human diseases^23, 24^. Except for one of the *HLA-C* alleles, all of the other alleles reached an 8-digit resolution, which is better than a recent prediction of HLA alleles based on whole-genome assemblies generated using the Oxford Nanopore data^25^ (Supplementary Table 2). We further assembled the two haplotypes using our 10x Genomics linked-read data from the targeted MHC region, and generated 18 contigs for each haplotype. For both haplotypes, the NGA50 scaffold size was consistently over 0.513 Mb with the largest contig length over 1.93 Mb (Supplementary Table 3).

**Table 1.** Genetic variants called from 10x Genomic linked-read data and the assembled haplotypes.

|  | 10x linked-read | Targeted assemblies | Garg *et al.*<sup>7</sup> |
| --- | --- | --- | --- |
| All variants | 28938 | 30100 | 30001 |
| Phased variants | 27244 | 30100 | 30001 |
| Heterozygous variants (GIAB) | 7534 (187) | 7608 (211) | 7616 (210) |
| Heterozygous variants (Illumina) | 14931 (2) | 15616 (0) | 15523 (8) |
| FNR (GIAB) | 0.02287 | 0.00513 | 0.00391 |
| FNR (Illumina) | 0.03599 | 0.01268 | 0.01679 |
| FPR (GIAB) | 0.61165 | 0.61948 | 0.61775 |
| FPR (Illumina) | 0.27994 | 0.29003 | 0.29063 |

The number of variants with different phasing information from the two benchmark datasets are indicated in the brackets. FNR: False negative rate; FPR: False positive rate.

Even though 10x Genomics linked-read sequencing performed well for phased variant calling, we were unable to achieve high-quality haplotype-resolved assemblies of the targeted MHC region. This result is consistent with recent reports showing that at least two sequencing platforms are needed to achieve high-quality haplotype-resolved *de novo* assemblies when trio information is not available^11, 17^. We therefore conducted additional sequencing using the PacBio HiFi platform. While the current deployment of the PacBio HiFi platform for genome assemblies requires large amounts of starting materials (e.g., more than 25 µg of genomic DNA from diploid human cells, in order to generate a minimum of 80 Gb data required for whole-genome assemblies), we were able to generate an average 30× coverage of the targeted MHC region by the 12 kb HiFi library (Supplementary Fig. 4a) from 20 ng of enriched HMW DNA. A highly reliable set of phased heterozygous variants was obtained by the intersection of variants generated from 10x Genomics linked-read data and HiFi reads to separate the HiFi reads into two haplotype-partitioned read sets. Eventually, two MHC haplotypes were *de novo* assembled separately from each haplotype-partitioned HiFi read set, together with untagged HiFi reads (Fig. 2a).

**Figure 2.**
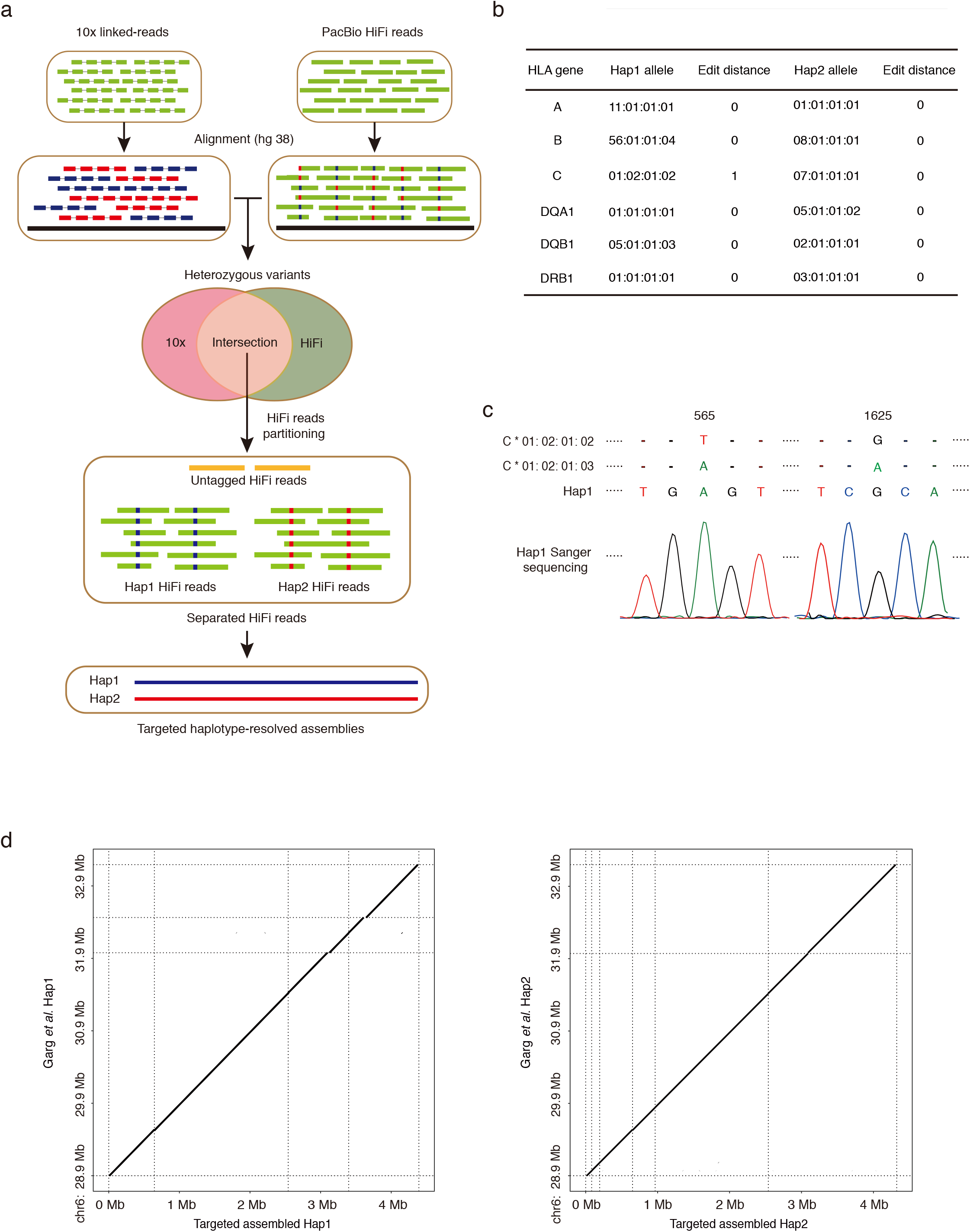
Targeted haplotype-resolved assemblies of the MHC region and HLA typing. **a**. Schematics of targeted haplotype-resolved assemblies of the MHC region with 10x linked-read data and PacBio HiFi reads. **b**. HLA typing of six classical *HLA* genes using the targeted haplotype-resolved assemblies. The edit distance is between our assemblies and the reported HLA alleles from the IMGT/HLA database. **c**. A novel *HLA-C* allele in the haplotype 1 is confirmed by Sanger sequencing. The nucleotides different from the best matching IMGT/HLA alleles are shown. The positions of the different nucleotides in the *HLA-C* gene are indicated. **d**. The comparison between our targeted assemblies and the MHC region from genome assemblies reported previously (Garg *et al*.)^7^ for each haplotype. The X-axis indicates the coordinate of our targeted assemblies, and the Y-axis indicates the coordinate of Garg *et al*.

The targeted assembled MHC haplotypes cover most of the MHC region of the hg38 reference (Table 2). We typed the 6 major *HLA* genes again, and all of them reached the 8-digit resolution (Fig. 2b). We noticed that the sequence of one *HLA-C* allele we identified is different from all the available *HLA-C* alleles reported in the IMGT/HLA database. Through Sanger sequencing, we confirmed that it is a novel *HLA-C* allele with one nucleotide difference from two best matching IMGT/HLA alleles (*C*01:02:01:02* and *C*01:02:01:03*) (Fig. 2c). These results showed that with our targeted haplotype-resolved assemblies of the MHC region, the HLA alleles can be typed with high accuracy.

**Table 2.** Statistics of two targeted assembled MHC haplotypes

|  | Haplotype 1 | Haplotype 2 |
| --- | --- | --- |
| **Aligned to the hg38 reference** |  |  |
| Fraction of the targeted region (%) | 98.368 | 95.847 |
| Total aligned length (bp) | 4311531 | 4194872 |
| NGA50 (bp) | 636872 | 456634 |
| LGA50 | 3 | 4 |
| **Without the reference** |  |  |
| Number of contigs | 4 | 6 |
| Largest contig length (bp) | 1895202 | 1784832 |
| Total length (bp) | 4391445 | 4323583 |

NGA50: minimum contig alignment length needed to cover 50% of the hg38 reference. LGA50: the number of contigs when reached NGA50.

### Evaluation of targeted haplotype-resolved assemblies of the MHC regio

Recently, a chromosome-scale high-quality haplotype-resolved assemblies of GM12878 (Garg *et al*. (2021))^7^ was reported using the PacBio HiFi and Hi-C data. To evaluate the performance of our targeted approach against the approach for genome assemblies, we compared our targeted haplotype-resolved assemblies of the MHC region to Garg *et al*., and observed a high consistency across the whole targeted region (Fig. 2d, Supplementary Fig. 4b). To evaluate the accuracy of our assembly result, variants were called by aligning the phased contigs against the hg38 reference, and then compared to the benchmarks of the GIAB (v3.3.2) and the Illumina Platinum Genomes. Compared to both benchmarks, we identified 30100 genetic variants within the targeted MHC region with FNRs lower than 1.3% (Table 1). All variants were phased with both switch error rates and Hamming error rates lower than 2% (Supplementary Table1), confirming the high quality of phasing. Notably, our targeted haplotype-resolved assemblies of the MHC region achieved a comparable consensus accuracy with Garg *et al*. (Table 1, Supplementary Table 1). Similar to Garg *et al*., we identified 61.9% and 29.0% new genetic variants compared to the GIAB (v.3.3.2) and the Illumina Platinum Genomes datasets, respectively (Table 1).

Based on the manual inspection of some of these newly identified variants, most of which are located in highly polymorphic or repetitive regions, and confirmed their accuracies by identifying supporting high-confidence HiFi reads (Fig. 3a).

**Figure 3.**
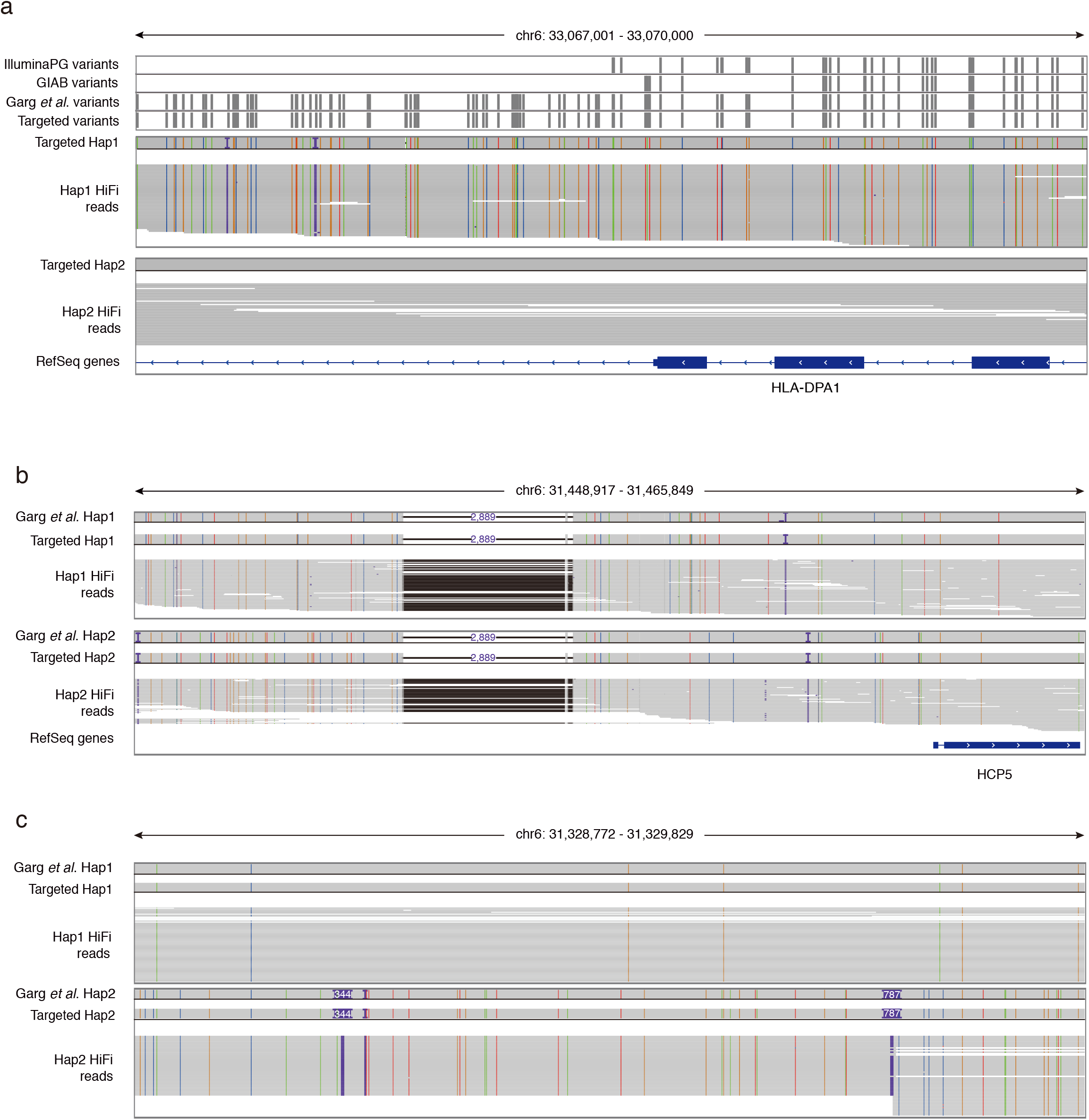
Accurate genetic variant calling using the targeted haplotype-resolved assemblies. **a**. An example of a region with newly identified variants. High-confidence HiFi reads with new variants are illustrated below. The colorful lines and dots positioned in the targeted assembly result of the haplotype 1 and HiFi reads indicate four types of nucleotides different from the hg38 reference. The purple character “I” and dots indicate small insertions (<50 bp) compared to the hg38 reference. Black dots indicate small deletions (<50 bp) compared to the hg38 reference. **b**. A 2,889 bp deletion upstream of the *HCP5* gene is illustrated and supported by HiFi reads for both haplotypes. **c**. A region with two insertions (344 bp and 787 bp) was identified and supported by HiFi reads.

With the targeted haplotype-resolved assemblies available, we were able to identify large insertions and deletions, which are difficult to find using short-read sequencing. For example, we identified a 2,889 bp deletion upstream of the *HCP5* gene (Fig. 3b) and a region with two insertions (344 bp and 787 bp) (Fig. 3c), both of which are supported by the presence of HiFi long reads and observed in Garg *et al*. It has been reported that the assembly-based approach enables more accurate and comprehensive characterization of genetic variants^1, 17^, and by comparing to the haplotype-resolved genome assemblies, our results showed that the targeted approach can achieve comparable completeness and accuracy on the detection of genetic variants in the targeted region.

### Accurate functional genomics analyses with the targeted haplotype-resolved assemblies

It has been reported that the most polymorphic parts of the MHC locus are located at regions around three HLA class I genes (*HLA-A, HLA-B*, and *HLA-C*) and three HLA class II genes (*HLA-DR, HLA-DQ*, and *HLA-DP*)^23, 26^. Consistently, our targeted haplotype-resolved assemblies showed that the major peaks of variants are in genomic regions around these six classical *HLA* genes (Supplementary Fig. 5a). While a high level of polymorphisms within the MHC region can be beneficial for a population facing an environment with various pathogens^27^, the presence of many polymorphisms and structural variants dramatically complicates the process of sequence alignment to a reference genome, and can generate ambiguous and even inaccurate results from short-read sequencing data^18, 28^. Consistent with this, using the hg38 as a reference for the sequence alignment of the RNA-Seq data generated from GM12878 cells, the number of sequencing reads mapped to the exons harboring many variants (regions highlighted with the gray shadow in Supplementary Fig. 5b) were much lower compared to the other exons from the same *HLA-B* gene. However, the effect of poor alignment was dramatically improved when we used the targeted assembled personal MHC haplotypes as the reference (Supplementary Fig. 5b). This highlights the importance of utilizing the targeted haplotype-resolved assemblies for accurate quantification of gene expression for genes with high polymorphisms.

This becomes more complicated in the study of DNA methylation. It is well-known that methylated cytosines are prone to spontaneous deamination, resulting the most common dinucleotide mutation (CG->TG) in mammalian genomes^29, 30^.

However, the gold-standard to quantify the level of DNA methylation at each CpG site after bisulfite-conversion is to calculate the number of reads sequenced with CG (indicating bisulfite-protected by the methyl group) divided by the number of reads sequenced with TG (indicating bisulfite-converted without the methyl group). In this situation, C/T variants on CpG sites will inevitably lead to erroneous methylation quantifications. For example, a specific TG-to-CG mutation was observed in the targeted assembled haplotype 2 positioned downstream of the *HLA-B* gene (the left grey box in Fig. 4a). For this position, if the sequencing reads were aligned to the hg38 reference, they will be missed from the evaluation of DNA methylation.

**Figure 4.**
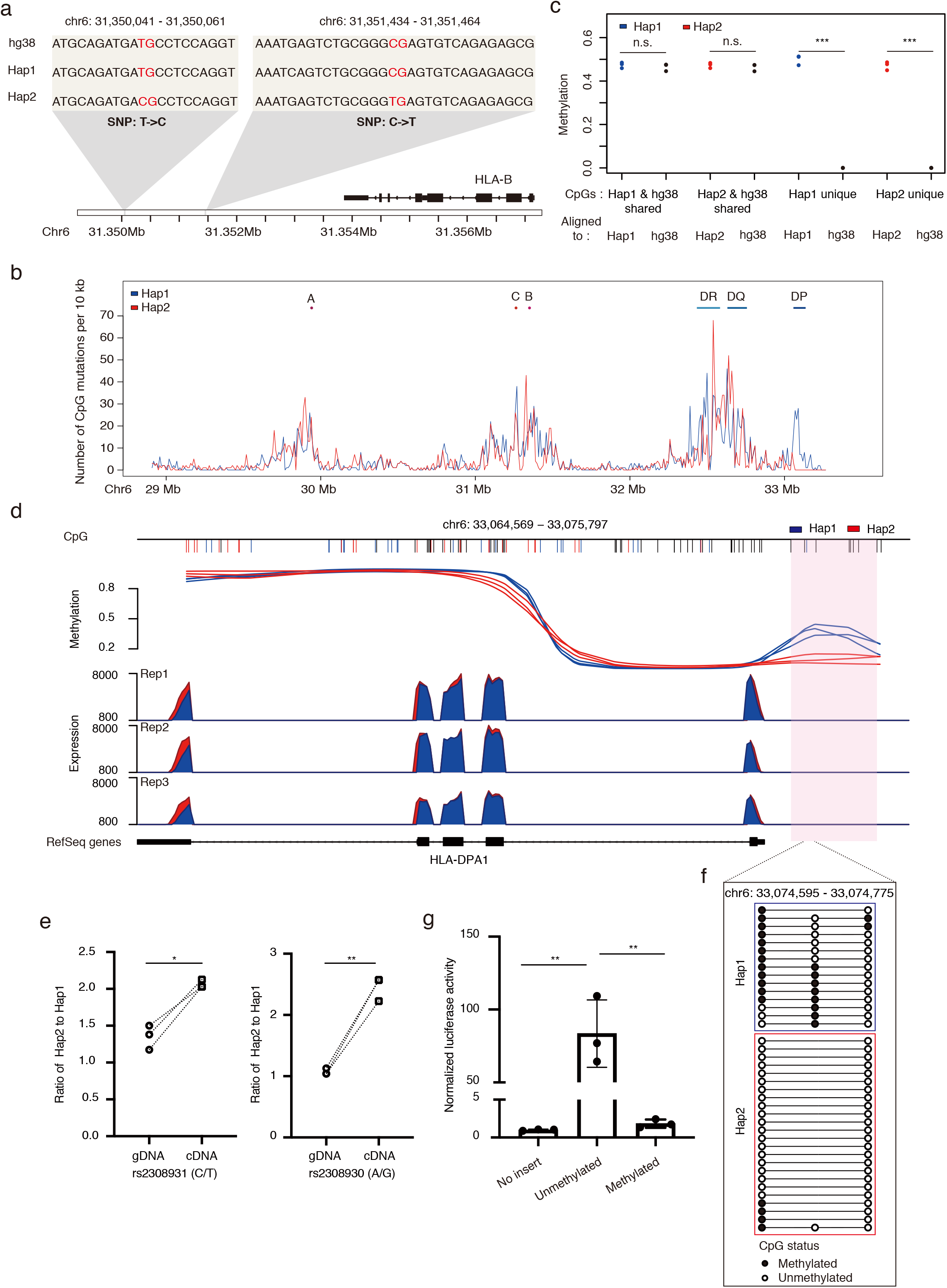
Accurate and allele-specific functional genomics analyses using the targeted haplotype-resolved assemblies. **a**. Examples of C/T variants on CpG sites. **b**. The CpG mutation density plot throughout the targeted MHC region. The number of CpG mutations is determined by the sum of CpG gains and losses relative to the hg38 reference per 10 kb bins. **c**. The effect of CpG mutations on DNA methylation analysis. Average DNA methylation levels for haplotype-specific or shared CpGs are estimated by aligning the sequencing reads to either corresponding targeted assembled personal MHC haplotype or the hg38 reference (\*\*\**P* value < 0.001, student’s t-test, two-sided). **d**. Allele-specific transcriptional regulation of the *HLA-DPA1* gene. Top part: allele-specific methylation in the promoter region of the *HLA-DPA1* gene. The differentially methylated region (DMR-DPA1) between two haplotypes is indicated as pink bar. The shared CpGs between two haplotypes are shown as black short lines, while haplotype-specific CpGs are indicated as blue or red short lines. The Y-axis indicates the DNA methylation level for each haplotype, separately. Three replicates are shown. Bottom part: allele-specific expression of the *HLA-DPA1* gene. The expression level for the haplotype 1 and 2 is indicated as blue and red, respectively. Three replicates are shown. **e**. Pyrosequencing of two polymorphic positions within the exons of the *HLA-DPA1* gene from either gDNA or cDNA of GM12878 cells. Y-axis indicates the ratio of the targeted SNPs between the two haplotypes. Data are represented from three independent experiments (\**P* value < 0.05, ** *P* value < 0.01, paired student’s t-test, two-sided). **f**. Bisulfite Sanger sequencing of PCR clones for part of the DMR-DPA1 region, containing one haplotype 1-specific CpG and two haplotype-shared CpGs. Each line represents one read where black or white circles illustrate methylated or unmethylated CpGs, respectively. **g**. DMR-DPA1 exerted methylation-dependent promoter activity on gene expression. Dual-luciferase reporter assays were performed using a CpG-free firefly luciferase reporter vector to evaluate the transcriptional activity of methylated or unmethylated DMR-DPA1 in HEK293T cells, with Firefly activity normalized to Renilla activity. Data are represented as mean ± SEM from three independent experiments (\*\**P* value < 0.01, student’s t-test, two-sided).

Alternatively, another CG-to-TG mutation observed in the haplotype 2 (the right grey box in Fig. 4a) will produce unmethylated calls without inducing any alignment mismatches and result in an excess of 0% methylated ‘sites’ at this position where there is actually no CpG site in the haplotype 2.

We evaluated the CpG sites among two targeted assembled MHC haplotypes and the hg38 reference, and found that 2.38% (1355/57010) of CpGs are specific in the haplotype 1 and 2.14% (1191/55659) are specific in the haplotype 2. However, 3.91% (2291/58570) of CpGs in the hg38 reference are not present in either haplotype (Supplementary Fig. 5c). When characterizing the distribution of these CpG mutations (haplotype-specific gains and losses of CpGs relative to the hg38) spanning the targeted MHC region, we observed that the major peaks are present in genomic regions around the six classical HLA genes (Fig. 4b), similar to the density plot generated from genetic variants (Supplementary Fig. 5a). The problematic effect resulting from CpG mutations was obvious when we performed the DNA methylation analyses separately for haplotype-specific or shared CpGs. As expected, extreme discrepancy was observed when computing the methylation level for haplotype-specific CpGs using the corresponding targeted assembled personal MHC haplotype or the hg38 as the reference, while this difference was not observed for shared CpGs (Fig. 4c). With the presence of the targeted haplotype-resolved assemblies of the MHC region, this bias can be easily addressed by aligning the bisulfite sequencing data to the targeted assembled personal MHC haplotypes as the reference, similar to what has been shown previously for analyzing methylation data on two highly divergent mouse strains^31^.

Bias for methylation quantification is also observed with DNA methylation arrays, reflecting that the high level of polymorphisms spanning the MHC region will affect the hybridization behavior of SNP-associated probes^32^. We observed a correlation coefficient (R^2^) of 0.96 for DNA methylation between the Illumina methylation EPIC beadchip and bisulfite sequencing data for CpGs without SNPs in the array probes (Supplementary Fig. 5d), while the correlation coefficient decreased to 0.9 for the probes harboring SNPs (Supplementary Fig. 5e). These results collectively highlight the utility of using the targeted assembled personal haplotypes as the reference for accurate quantifications of DNA methylation and gene expression data, especially for highly polymorphic regions such as the MHC.

### Allele-specific analyses with the targeted haplotype-resolved assemblies

With the targeted assembled MHC haplotypes available, we further characterized allele-specific methylation and expression for diploid GM12878 cells. We first analyzed the allele-specific expression (ASE) using two targeted assembled personal MHC haplotypes as the reference, and identified that seven genes within the targeted MHC region exhibit allele-specific expression (adjusted P-value ≤ 0.05) (Supplementary Fig. 6a and Supplementary Table 4). For the expression of the *HLA-DPA1* gene, which had a 1.1-fold difference between the two haplotypes, we observed consistently higher expression for all of the haplotype 2 exons of *DPA1* (Fig. 4d), and this ASE was validated using pyrosequencing (Fig. 4e). The fact that the expression of the *DPA1* gene in the haplotype 2 (*DPA1*01:03:01:02*) is higher than that in the haplotype 1 (*DPA1*02:01:01:02*) in GM12878 cells is consistent with a recent finding that the expression of the *DPA1*01* allele is significantly higher than that of the *DPA1*02* allele in the population^33^.

We then analyzed the allele-specific methylation (ASM) for the targeted MHC region, and found 211 differentially methylated regions (DMRs) between the two haplotypes (Supplementary Table 5). We included haplotype-specific CpGs in the methylation analysis, as this will increase the accuracy and power in the differential analysis^31^ (Supplementary Fig. 6b). We noticed that, in the promoter region of the *HLA-DRA1* gene, there was a differentially methylated region (termed DMR-DPA1 hereafter), in which the haplotype 1 was hypermethylated (Fig. 4d). DMR-DPA1 contains one haplotype 1-specific CpG site and several shared CpGs, and the allele-specific methylation of DMR-DPA1 was validated using bisulfite Sanger sequencing (Fig. 4f).

To examine whether the ASM of DMR-DPA1 exerts any regulatory impacts on the ASE of the *DPA1* gene, we performed a cell-based dual luciferase reporter assay and detected a significant increase of gene expression when unmethylated DMR-DPA1 is present. In contrast, this upregulation of gene expression is abolished when DMR-DPA1 on the luciferase reporter was methylated *in vitro* (Fig. 4g). This suggests that the ASE of the *HLA-DPA1* gene can be negatively regulated through ASM on the promoter region of *HLA-DPA1*. These results illustrate that the targeted assembled personal haplotypes enable allele-specific functional genomics analyses in diploid human cells, and can contribute to our understandings of allele-specific transcriptional regulation of targeted genomic regions.

## Discussion

The recent advances of constructing high-quality haplotype-resolved genome assemblies have made it possible to comprehensively discover genetic variants at the chromosome-scale. However, this genome-wide approach is difficult to be applied in a large group of individuals or even among diverse populations to fully characterize genetic variants in order to reveal their potential functional relevance. In this study, using the clinically important MHC region as an example, by combining the CRISPR-based enrichment with 10x Genomics linked-read sequencing and the PacBio HiFi platform, we successfully *de novo* assembled the targeted MHC haplotypes without parent-child trio information. By comparing to the haplotype-resolved genome assemblies from the same cell line, we showed that our targeted approach for obtaining haplotype-resolved assemblies achieved comparable completeness and accuracy with greatly reduced computing complexity, sequencing cost, as well as the amount of starting materials.

Even though genome-wide association studies (GWAS) have greatly advanced our understanding on the human genome, it is still challenging to pinpoint causal variants involved in disease etiology for genetic regions with high polymorphisms and strong linkage disequilibrium (e.g., the MHC region). It has been suggested that haplotype-based approaches may overcome these challenges towards more accurate genome inference^18^. One example for this is the haplotypes of HLA-DRB1-HLA-DQA1-HLA-DQB1 greatly increased the risk of type 1 diabetes^34^. Another example is the identification of different alleles of the complement component 4 (C4) genes within the MHC region for the development of schizophrenia^35^. Considering the MHC region is one of the most difficult regions to be assembled in human genome^17^, our targeted approach can (in theory) be adopted for haplotype-based population genetic studies to analyze any genomic regions of interest.

In addition to genetic variants, functional analyses of any genomic regions can also benefit greatly from our unbiased approach of constructing haplotype-resolved assemblies of targeted regions. For regions with high polymorphisms, such as the MHC, the conventional strategy of mapping short-read sequencing data to a single reference genome (e.g., hg19 or hg38) is known to yield biased alignments, leading towards inaccurate quantifications^36, 37^. This has prompted recent attempts to replace a single reference genome with the computationally inferred personal genotypes as the reference, all of which showed improved accuracies of quantifications compared to the standard approach^33, 38-41^. However, these computational strategies highly depend on the completeness of our understanding on diverse human genomes, which is still far from saturation, even for the widely studied classical *HLA* genes^42^. Here, using the targeted assembled personal MHC haplotypes as an example, we not only showed that the results of functional genomics analyses of the MHC region are much more accurate, but also revealed allele-specific transcriptional regulation of the *HLA-DPA1* gene. It can be safely assumed that other functional genomics analyses of any targeted genomic regions would benefit from our approach of using haplotype-resolved assemblies of targeted regions as the personal reference. This integrated analysis of genetic and epigenetic data may pave a new way for comprehensively elucidating the molecular mechanisms in disease etiology.

## Acknowledgements

This work was supported by the grants from the National Natural Science Foundation of China (No. 82171837 to YL), the National Key R&D Program of China (No. 2021YFC2701001 to YL), Program of Shanghai Academic/Technology Research Leader Grant (No. 20XD1420400 to YL), Shanghai Municipal Science and Technology Major Project (No. 2017SHZDZX01 and No. 2018SHZDZX01 to YL) and ZJLab, and the Shanghai Science and Technology Foundation (20ZR1404700 to YL). We thank Qianjie Wang and Wilson Cheng from PacBio APAC for useful discussion, troubleshooting efforts and analysis suggestions.

