## Supplementary Figures for "CRISPR-based targeted haplotype-resolved assemblies of a megabase region"

**Supplementary Figure 1. The CRISPR-based targeted enrichment of a megabase region**

**a**. Schematics of the CRISPR-based targeted enrichment. Cells were embedded in agarose plugs and digested with proteinase K. The targeted genomic region was cleaved by CRISPR-based in-gel digestion and separated by PFGE. The final targeted HMW MHC molecules were recovered through dialysis. **b**. The cleavage efficiencies of the designed sgRNAs were evaluated individually. The efficiency was evaluated through *in vitro* cleavage assays with PCR products amplified from the targeted regions. Two sets of sgRNAs chosen for the final targeted enrichment of the MHC region are highlighted with red and blue texts, respectively.

**Supplementary Figure 2. Targeted separation and recovery of the HMW MHC molecules**

The targeted enriched HMW DNA molecules were still longer than 50 kb indicated by the CHEF DNA size standards-Lambda Ladder (M).

**Supplementary Figure 3. Schematics of phased variant calling and HLA typing with 10x Genomics linked-read data**

The 10x Genomics linked-read data was aligned to the hg38 reference using Long Ranger v.2.2.2 to call phased variants within the targeted MHC region. The accuracy of phased variants was evaluated by comparing to the GIAB v3.3.2 and the Illumina Platinum Genomes datasets. The 10x Genomics linked reads were split into two haplotype-partitioned read sets based on their phasing information generated from Long Ranger, and HLA types for each haplotype were predicted separately using the

HLA-VBseq (PMID: 31240105) with the IMGT/HLA database (v3.44).

**Supplementary Figure 4. Targeted haplotype-resolved assemblies of the MHC region**

**a**. The coverage of the PacBio HiFi reads on the targeted MHC region is shown. The digestion positions for two sets of sgRNAs are indicated as red and blue bars, respectively. **b**. The comparison of our targeted assemblies and genome assemblies reported previously (Garg *et al.*) (PMID: 33288905) shows high consistency across the targeted MHC region for each haplotype. The consistent assemblies are indicated as gray lines. The different colored horizontal bars represent contigs.

**Supplementary Figure 5. Functional genomics analyses with the targeted haplotype-resolved personal assemblies**

**a.** The density plot of genetic variants throughout the targeted MHC region. The X-axis indicates the coordinates of the targeted MHC assemblies and the Y-axis indicates the number of genetic variants (SNPs and InDels) relative to hg38 reference in each 10 kb window. Blue line: the assembly of haplotype 1; red line: the assembly of haplotype 2. Red dots: locations of three classical HLA I genes; blue bars: locations of three classical HLA II genes. **b**. Quantification of RNA-Seq reads on the HLA-B gene based on difference references (indicated at the left). Three replicates are shown. Two regions with high density of SNPs which have biased sequence alignment are indicated as grey bars. **c**. The Venn diagram shows the numbers of CpGs specific or shared by the assembly of each haplotype and the hg38 reference. **d** and **e**. The methylation correlations between the EPIC array and bisulfite sequencing data. We analyzed the methylation level for shared CpGs between the Illumina Infinium EPIC methylation array and bisulfite sequencing data. The shared CpGs were separated into two groups by whether genetic variants are present within 5 bp of the interrogated CpG site on the probes of EPIC array (**d**), or absent from the probes (**e**).

**Supplementary Figure 5. Analyses of allele-specific expression and methylation**

**a**. The volcano plot of genes that are differentially expressed between two alleles. The red line represents the cut-off for statistical significance of adjusted P-value less than 0.05. Red dots indicate the allele-specific expressed genes. **b**. The allele-specific methylation analysis with (the top panel) or without (the bottom panel) the haplotype-specific CpGs included. The methylation level of the promoter region of *HCG18* gene is shown as an example. The identified DMR indicating allele-specific methylation is indicated with the pink box. The shared CpGs between two haplotypes are shown as black short lines, while haplotype-specific CpGs are indicated as blue or red short lines.

### Supplementary Figure. 1

a

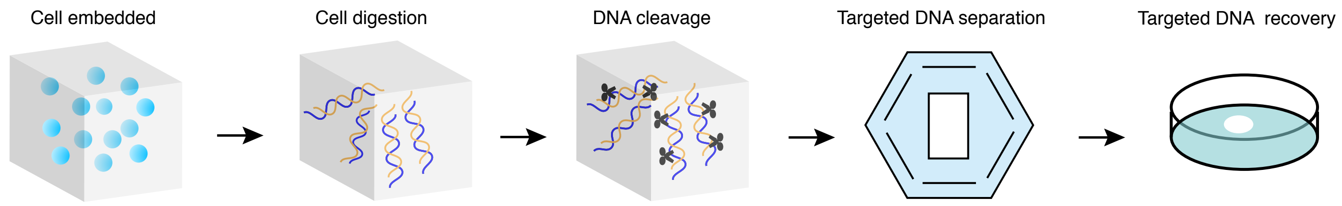


b

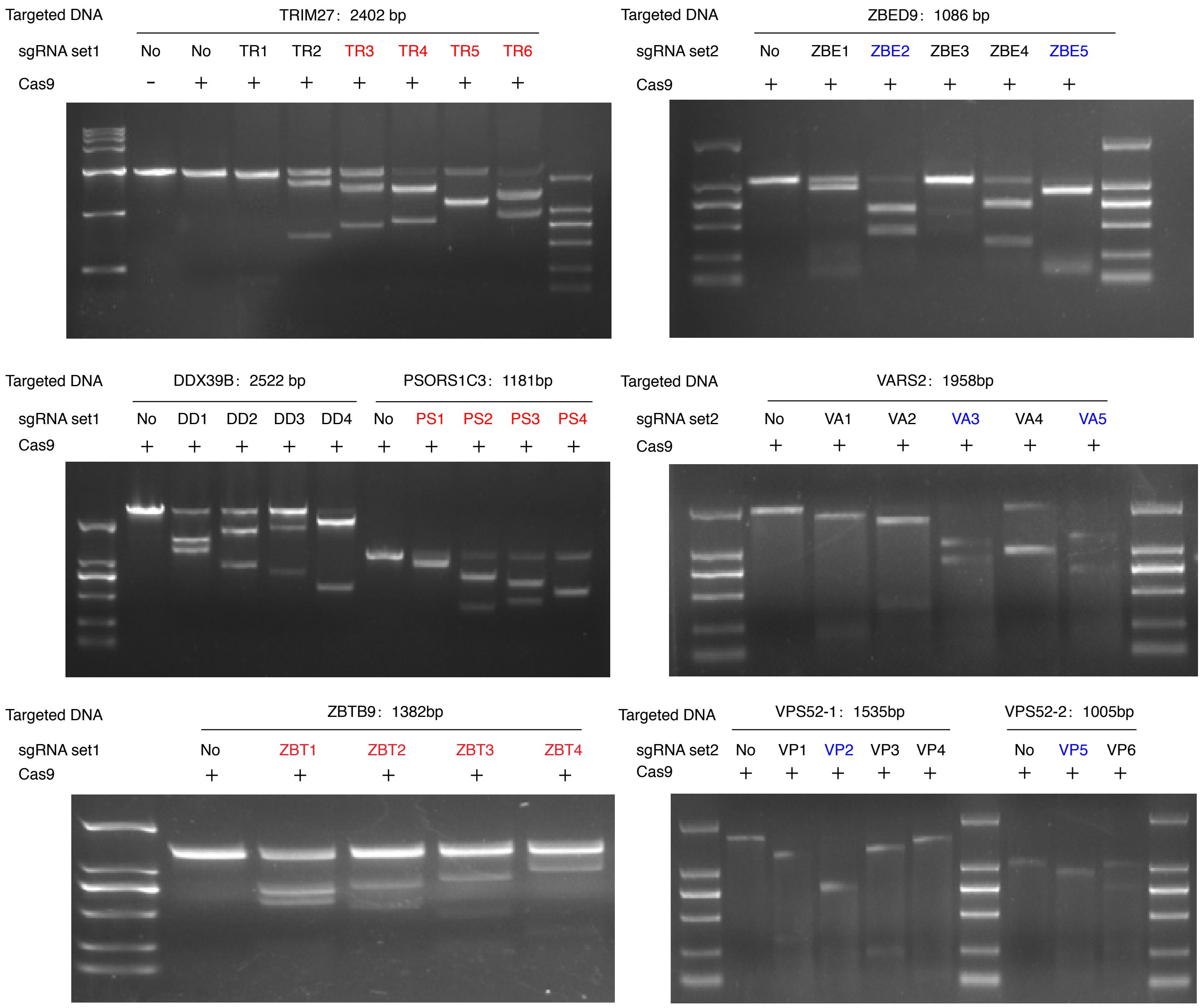

#### Supplementary Figure. 2

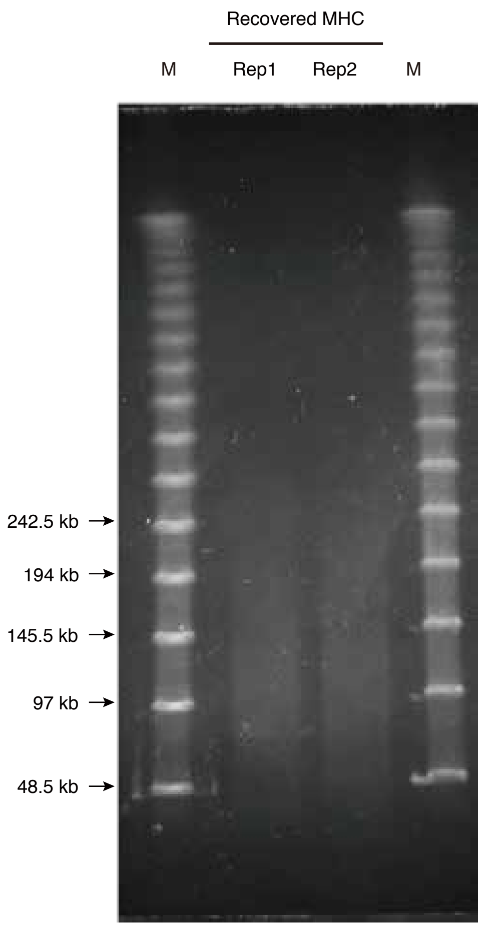

#### Supplementary Figure. 3

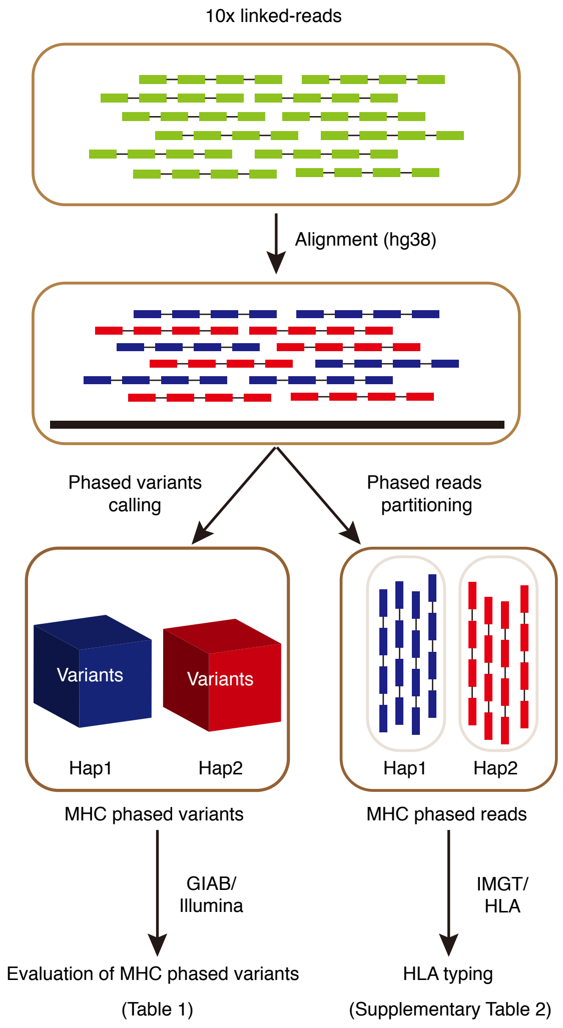

#### Supplementary Figure. 4

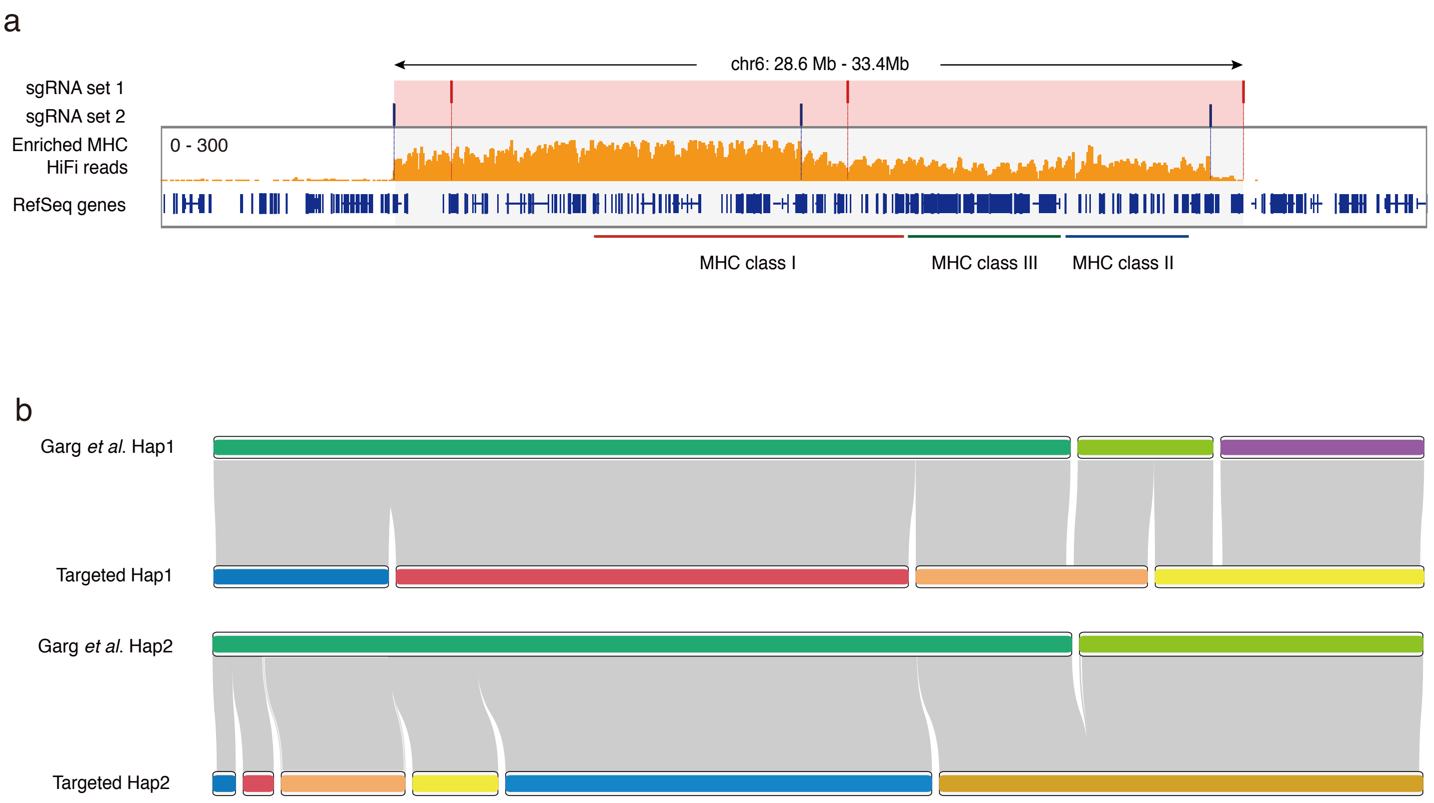

#### Supplementary Figure. 5

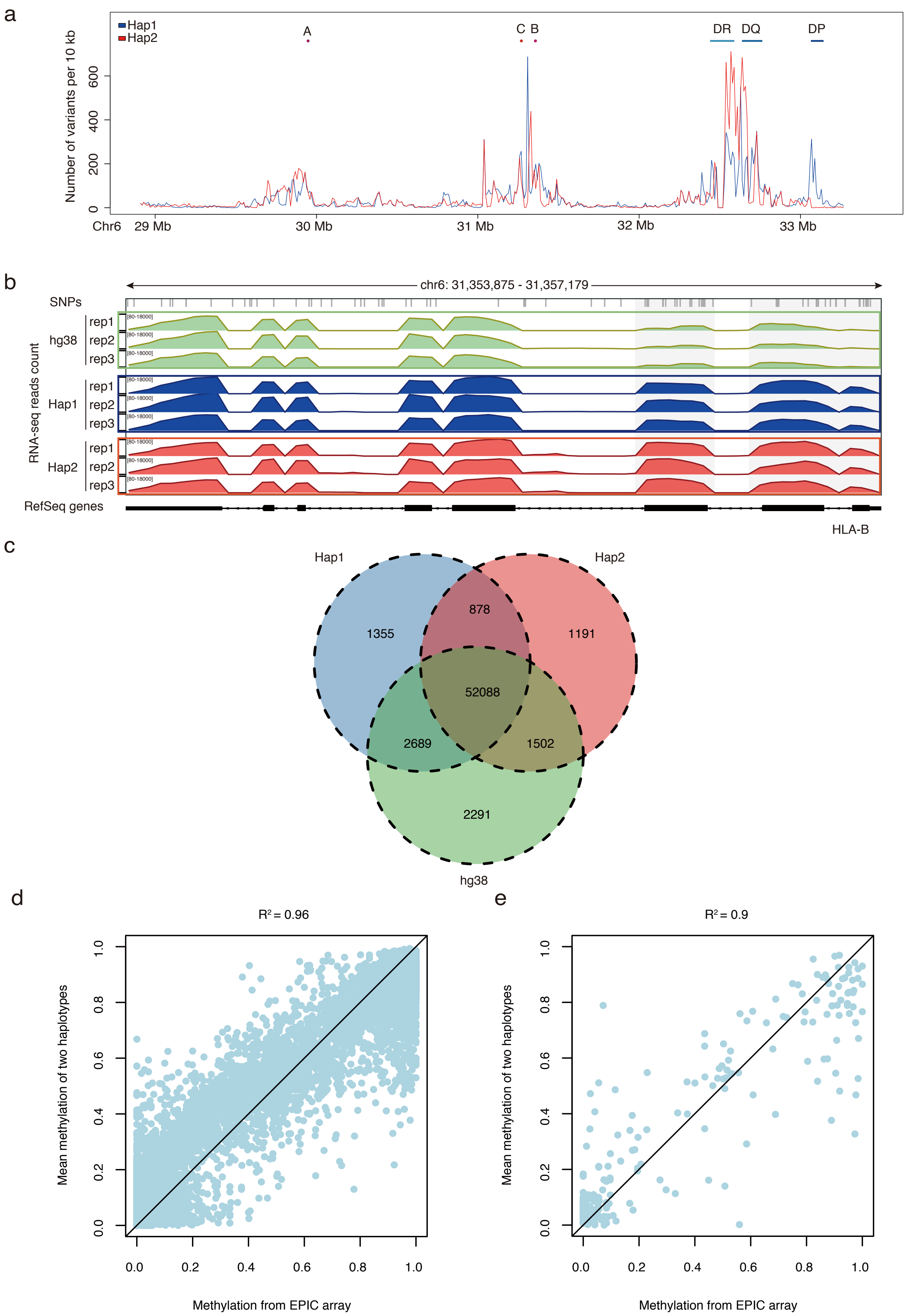

#### Supplementary Figure. 6

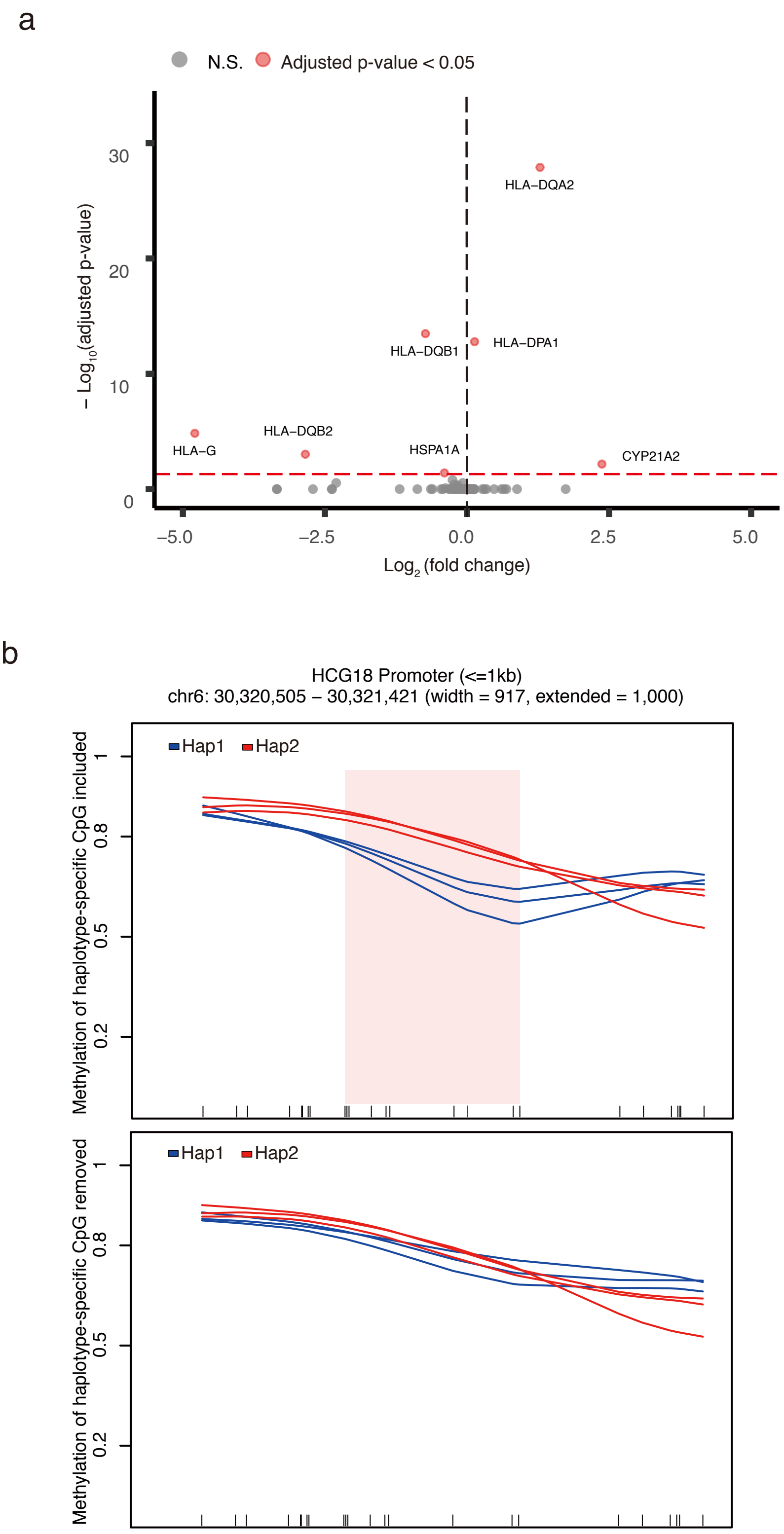
