## Supplementary Tables for "CRISPR-based targeted haplotype-resolved assemblies of a megabase region"

**Supplementary Table 1. The performance of phased variant calling**

| | Homo-variants (GIAB) | Homo-variants (Illumina) | Switch error (GIAB) | Switch error (Illumina) | Hamming error (GIAB) | Hamming error (Illumina) |
|---|---|---|---|---|---|---|
| 10x linked-reads | 3444 | 5966 | 1.66% | 0.42% | 1.48% | 0.31% |
| Garg *et al.* | 3483 | 5996 | 1.75% | 0.43% | 1.76% | 0.36% |
| Targeted assemblies | 3476 | 6028 | 1.75% | 0.42% | 1.77% | 0.31% |

Switch error: the percentage of adjacent SNP pairs wrongly phased in comparison to the benchmarks

Hamming error: the percentage of SNPs wrongly phased in comparison to the benchmarks

**Supplementary Table 2. HLA typing using 10x Genomics linked-read data**

| | Alleles typed using 10x Genomics linked-read data | | Alleles predicted by Jain *et al.* | |
|---|---|---|---|---|
| | Hap1 | Hap2 | Hap1 | Hap2 |
| HLA-A | 11:01:01:01 | 01:01:01:01 | 11:01:01G | 01:01:01G |
| HLA-B | 56:01:01:04 | 08:01:01:01 | 56:01:01G | 08:01:01G |
| HLA-C | 01:148 (...) | 07:01:01:01 | 01:02:01G | 07:01:01G |
| HLA-DQA1 | 01:01:01:01 | 05:01:01:02 | 01:01:01G | 05:01:01G |
| HLA-DQB1 | 05:01:01:03 | 02:01:01:01 | 05:01:01G | 02:01:01G |
| HLA-DRB1 | 01:01:01:01 | 03:01:01:01 | 01:01:01G | 03:01:01G |

HLA typing for six classical HLA genes.
For the haplotype 1 of the *HLA-C* gene, several alleles were typed.
Information regarding the ‘G’ group in HLA alleles predicted by Jain *et al.*
is available at ‘http://hla.alleles.org/alleles/g groups.html’

### Supplementary Table 3. Assembly statistics of the targeted MHC region from 10x Genomics linked-read data

| | Hap1 | Hap2 |
|---|---|---|
| **Aligned to the hg38 reference** | | |
| Fraction of the targeted region (%) | 93.637 | 93.8 |
| Total aligned length (bp) | 4153251 | 4159618 |
| NGA50 (bp) | 513265 | 513209 |
| LGA50 | 3 | 3 |
| **Without the reference** | | |
| Number of contigs | 18 | 18 |
| Largest contig length (bp) | 1930349 | 1936502 |
| Total length (bp) | 4267591 | 4274584 |

The assembly statistics of the targeted MHC region were calculated by comparing to the hg38 reference using alternate contigs over 10 kb

**Supplementary Table 4. The genes with allele-specific expression**

| | Chromosome | Start (hg38) | End (hg38) | Fold Change | log2FoldChange | P-value | Adjusted p-value |
|---|---|---|---|---|---|---|---|
| *HLA-G* | chr6 | 29826979 | 29831122 | 0.03615182 | -4.789788082 | 2.97E-07 | 1.45E-05 |
| *HSPA1A* | chr6 | 31815514 | 31817942 | 0.75898766 | -0.397851663 | 0.00142415 | 0.03967281 |
| *CYP21A2* | chr6 | 32035476 | 32041670 | 5.18405875 | 2.374082067 | 0.00020719 | 0.006733791 |
| *HLA-DQB1* | chr6 | 32659464 | 32666689 | 0.60212056 | -0.731875725 | 3.39E-16 | 3.31E-14 |
| *HLA-DQA2* | chr6 | 32741386 | 32746887 | 2.43726332 | 1.28526213 | 6.53E-31 | 1.27E-28 |
| *HLA-DQB2* | chr6 | 32756098 | 32763553 | 0.1391563 | -2.845221866 | 2.42E-05 | 0.000942132 |
| *HLA-DPA1* | chr6 | 33064569 | 33080778 | 1.09691986 | 0.133458123 | 2.55E-15 | 1.66E-13 |

**Supplementary Table 5. The allele-specific DNA methylation regions within the targeted MHC region**

| Chr. | Start (hg38) | End (hg38) | Number of CpGs | Width (bp) | Area T-Stat | Mean methylation in Hap1 | Mean methylation in Hap2 | Methylation difference |
|---|---|---|---|---|---|---|---|---|
| chr6 | 32013394 | 32019492 | 87 | 6099 | -1333.81847 | 0.014356404 | 1 | -0.985643596 |
| chr6 | 32007839 | 32011194 | 135 | 3356 | -913.525948 | 0.356441166 | 1 | -0.643558834 |
| chr6 | 31498441 | 31499244 | 48 | 804 | -625.191895 | 0.270104949 | 0.711866343 | -0.441761394 |
| chr6 | 32602234 | 32604549 | 41 | 2316 | 578.564661 | 0.724213496 | 0.182885762 | 0.541327733 |
| chr6 | 32445453 | 32449616 | 56 | 4164 | -575.266799 | 0.616907209 | 0.800641902 | -0.183734693 |
| chr6 | 31307887 | 31309034 | 74 | 1148 | 562.9821133 | 0.562750474 | 0.125237314 | 0.43751316 |
| chr6 | 32653424 | 32656442 | 51 | 3019 | 548.5146107 | 0.899470704 | 0.511228028 | 0.388242676 |
| chr6 | 31491215 | 31492034 | 29 | 820 | 530.161043 | 0.713720279 | 0.263256758 | 0.45046352 |
| chr6 | 32384231 | 32385301 | 65 | 1071 | -474.046886 | 0.570610511 | 0.735644061 | -0.16503355 |
| chr6 | 29015123 | 29015920 | 52 | 798 | 470.7000728 | 0.575131226 | 0.271396782 | 0.303734444 |
| chr6 | 32459689 | 32460769 | 42 | 1081 | -439.590722 | 0.070631337 | 0.242207268 | -0.171575931 |
| chr6 | 31185055 | 31186666 | 35 | 1612 | -433.631882 | 0.189794311 | 0.556077987 | -0.366283676 |
| chr6 | 32632514 | 32636619 | 47 | 4106 | 424.6216042 | 0.642693857 | 0.247190796 | 0.395503061 |
| chr6 | 32024010 | 32026432 | 72 | 2423 | 412.4889843 | 0.796822174 | 0.46480401 | 0.332018164 |
| chr6 | 32605330 | 32607301 | 34 | 1972 | -364.973763 | 0.639168901 | 0.896199241 | -0.257030339 |
| chr6 | 31766033 | 31766803 | 37 | 771 | -327.523267 | 0.525666688 | 0.737408973 | -0.211742285 |
| chr6 | 33125332 | 33126335 | 28 | 1004 | -299.285534 | 0.344708616 | 0.733075736 | -0.38836712 |
| chr6 | 29722249 | 29722516 | 24 | 268 | -299.1679 | 0.354606804 | 0.862222346 | -0.507615542 |
| chr6 | 32599095 | 32601653 | 46 | 2559 | -280.647601 | 0.751587575 | 0.857293412 | -0.105705838 |
| chr6 | 29972074 | 29974929 | 28 | 2856 | 254.9386577 | 0.651302063 | 0.357107396 | 0.294194668 |
| chr6 | 31455702 | 31456321 | 46 | 620 | 254.0261533 | 0.631361309 | 0.228541373 | 0.402819936 |
| chr6 | 32641279 | 32643872 | 32 | 2594 | -239.823018 | 0.59014771 | 0.743300351 | -0.153152641 |
| chr6 | 31357727 | 31358694 | 13 | 968 | -236.433283 | 0.154651112 | 0.645255737 | -0.490604625 |
| chr6 | 32457117 | 32459429 | 29 | 2313 | 236.154491 | 0.670794749 | 0.367220906 | 0.303573843 |
| chr6 | 30906938 | 30907680 | 23 | 743 | 234.9769015 | 0.809011255 | 0.576159679 | 0.232851576 |
| chr6 | 32610659 | 32612490 | 21 | 1832 | 224.0525197 | 0.744992281 | 0.448071002 | 0.296921279 |
| chr6 | 32552141 | 32555128 | 36 | 2988 | -206.69207 | 0.300466107 | 0.585654005 | -0.285187899 |
| chr6 | 29925602 | 29926420 | 28 | 819 | 206.1842956 | 0.882005924 | 0.642732912 | 0.239273012 |

| | | | | | | | | |
|---|---|---|---|---|---|---|---|---|
| chr6 | 32689765 | 32692597 | 23 | 2833 | 203.2840903 | 0.391013067 | 0.129784726 | 0.261228341 |
| chr6 | 33104744 | 33109156 | 29 | 4413 | -195.543912 | 0.433768203 | 0.997528281 | -0.563760078 |
| chr6 | 31495843 | 31497203 | 25 | 1361 | 193.1517236 | 0.519316794 | 0.172078913 | 0.347237881 |
| chr6 | 31198592 | 31199650 | 20 | 1059 | 191.9819196 | 0.423960857 | 0.20242015 | 0.221540707 |
| chr6 | 29921260 | 29925175 | 35 | 3916 | -190.81155 | 0.676017082 | 0.844407784 | -0.168390702 |
| chr6 | 32547872 | 32548687 | 36 | 816 | 189.5560514 | 0.743565831 | 0.558996442 | 0.18456939 |
| chr6 | 32607736 | 32609307 | 26 | 1572 | 189.1355258 | 0.556244945 | 0.310798741 | 0.245446204 |
| chr6 | 32699402 | 32700654 | 32 | 1253 | -188.352431 | 0.348763686 | 0.458911657 | -0.110147972 |
| chr6 | 32450379 | 32453449 | 32 | 3071 | 180.3450864 | 0.517675817 | 0.275399518 | 0.242276299 |
| chr6 | 33101196 | 33102887 | 13 | 1692 | 166.2273507 | 0.398294858 | 0.221429448 | 0.176865411 |
| chr6 | 29680707 | 29681184 | 37 | 478 | 159.2938876 | 0.57519346 | 0.405993488 | 0.169199971 |
| chr6 | 29633557 | 29634427 | 17 | 871 | -159.141185 | 0.195450752 | 0.385117066 | -0.189666314 |
| chr6 | 30813662 | 30814894 | 19 | 1233 | 157.7922264 | 0.724450202 | 0.45285196 | 0.271598242 |
| chr6 | 32237504 | 32239684 | 26 | 2181 | 155.2853162 | 0.570198532 | 0.446468721 | 0.123729811 |
| chr6 | 31350866 | 31352598 | 24 | 1733 | 154.9224637 | 0.726553951 | 0.496126667 | 0.230427284 |
| chr6 | 32473210 | 32474715 | 24 | 1506 | 154.7751544 | 0.614114332 | 0.415812695 | 0.198301638 |
| chr6 | 31054591 | 31056091 | 23 | 1501 | 152.8894258 | 0.427069711 | 0.246696827 | 0.180372884 |
| chr6 | 32761768 | 32762143 | 35 | 376 | -152.516274 | 0.28803522 | 0.448002304 | -0.159967084 |
| chr6 | 31981088 | 31982887 | 31 | 1800 | 150.3999435 | 0.82118051 | 0.449644081 | 0.37153643 |
| chr6 | 31187135 | 31188369 | 24 | 1235 | -144.885962 | 0.596783066 | 0.792932669 | -0.196149603 |
| chr6 | 32938463 | 32939608 | 15 | 1146 | -141.168129 | 0.315228578 | 0.568060382 | -0.252831804 |
| chr6 | 31309393 | 31311231 | 23 | 1839 | -137.979577 | 0.178055086 | 0.685982034 | -0.507926947 |
| chr6 | 32737471 | 32738699 | 22 | 1229 | -137.441264 | 0.382287797 | 0.619040946 | -0.236753149 |
| chr6 | 31266724 | 31268273 | 24 | 1550 | -136.993965 | 0.622959303 | 0.820575799 | -0.197616496 |
| chr6 | 29679798 | 29680384 | 24 | 587 | -128.784759 | 0.497943577 | 0.678713519 | -0.180769942 |
| chr6 | 30801437 | 30802643 | 26 | 1207 | 127.330797 | 0.840764488 | 0.727442769 | 0.113321719 |
| chr6 | 31274144 | 31275826 | 15 | 1683 | 126.9568052 | 0.798666847 | 0.617338301 | 0.181328546 |
| chr6 | 29749411 | 29750140 | 14 | 730 | 125.9695318 | 0.604406839 | 0.374519714 | 0.229887126 |
| chr6 | 29850004 | 29850377 | 26 | 374 | -121.189769 | 0.253947753 | 0.387013036 | -0.133065283 |
| chr6 | 31080529 | 31081559 | 16 | 1031 | -118.491765 | 0.445827108 | 0.600150818 | -0.154323709 |
| chr6 | 32543683 | 32544680 | 15 | 998 | 114.0975723 | 0.592659052 | 0.390158931 | 0.202500121 |
| chr6 | 32391946 | 32393205 | 15 | 1260 | 112.1989657 | 0.631733109 | 0.336595751 | 0.295137358 |

| | | | | | | | | |
|---|---|---|---|---|---|---|---|---|
| chr6 | 33123974 | 33124773 | 18 | 800 | -109.848494 | 0.465846683 | 0.690199848 | -0.224353165 |
| chr6 | 28976238 | 28977177 | 21 | 940 | 108.764235 | 0.774002337 | 0.668655178 | 0.10534716 |
| chr6 | 31433747 | 31436078 | 23 | 2332 | -105.556723 | 0.22791718 | 0.336682368 | -0.108765188 |
| chr6 | 31333881 | 31335616 | 21 | 1736 | -103.935708 | 0.213706998 | 0.49467826 | -0.280971263 |
| chr6 | 33109854 | 33112277 | 15 | 2424 | -101.805148 | 0.456134343 | 0.905046719 | -0.448912376 |
| chr6 | 31279188 | 31280656 | 22 | 1469 | -101.699693 | 0.398274725 | 0.526660526 | -0.1283858 |
| chr6 | 30809460 | 30810400 | 17 | 941 | -100.126852 | 0.537170779 | 0.697124581 | -0.159953802 |
| chr6 | 32530092 | 32531364 | 10 | 1273 | -98.2855882 | 0.394486475 | 0.878284978 | -0.483798503 |
| chr6 | 29803011 | 29805278 | 15 | 2268 | -98.1463215 | 0.073681917 | 0.259919934 | -0.186238018 |
| chr6 | 31114410 | 31115545 | 19 | 1136 | 97.0272102 | 0.272060289 | 0.111928525 | 0.160131764 |
| chr6 | 31485986 | 31486502 | 12 | 517 | 96.8118836 | 0.879965182 | 0.554135124 | 0.325830059 |
| chr6 | 29886854 | 29887495 | 12 | 642 | 96.10461988 | 0.4437775 | 0.194569015 | 0.249208485 |
| chr6 | 32559811 | 32561666 | 22 | 1856 | 94.99828674 | 0.571936773 | 0.353479689 | 0.218457083 |
| chr6 | 31542914 | 31543237 | 12 | 324 | -94.2910875 | 0.218259625 | 0.382499233 | -0.164239609 |
| chr6 | 32442500 | 32443598 | 19 | 1099 | 93.62710473 | 0.693198521 | 0.507275458 | 0.185923063 |
| chr6 | 29638884 | 29639651 | 19 | 768 | 93.354658 | 0.429028202 | 0.297956989 | 0.131071213 |
| chr6 | 32258703 | 32260485 | 14 | 1783 | 93.04209716 | 0.502135436 | 0.344211129 | 0.157924307 |
| chr6 | 29888586 | 29888787 | 22 | 202 | -92.8103742 | 0.619582112 | 0.763477399 | -0.143895287 |
| chr6 | 32484741 | 32486210 | 18 | 1470 | -91.7594618 | 0.431780138 | 0.811503196 | -0.379723058 |
| chr6 | 31272647 | 31273359 | 11 | 713 | -91.287611 | 0.097742548 | 0.406795928 | -0.30905338 |
| chr6 | 31296744 | 31298312 | 15 | 1569 | -88.7571587 | 0.15241174 | 0.345707029 | -0.193295289 |
| chr6 | 32664223 | 32664497 | 14 | 275 | 87.08677824 | 0.385436864 | 0.032253744 | 0.353183119 |
| chr6 | 31127499 | 31127878 | 16 | 380 | 85.81381999 | 0.380257423 | 0.190763037 | 0.189494386 |
| chr6 | 31632918 | 31633752 | 18 | 835 | 85.76412429 | 0.890568994 | 0.77695723 | 0.113611764 |
| chr6 | 31270684 | 31271173 | 21 | 490 | 85.7408015 | 0.524450268 | 0.127856091 | 0.396594178 |
| chr6 | 31001118 | 31001843 | 18 | 726 | 83.33184308 | 0.477496576 | 0.352554846 | 0.124941731 |
| chr6 | 33061499 | 33062484 | 8 | 986 | 82.23395793 | 0.876355789 | 0.491524892 | 0.384830896 |
| chr6 | 30102179 | 30102332 | 18 | 154 | 82.05408611 | 0.2831283 | 0.173175691 | 0.109952609 |
| chr6 | 32414843 | 32415404 | 11 | 562 | -80.9910158 | 0.21478218 | 0.4176921 | -0.20290992 |
| chr6 | 29724693 | 29725757 | 18 | 1065 | 79.63684394 | 0.796595404 | 0.644855601 | 0.151739803 |
| chr6 | 29933133 | 29933674 | 14 | 542 | -77.4043074 | 0.647350025 | 0.860747043 | -0.213397018 |
| chr6 | 32798159 | 32798767 | 12 | 609 | -77.2256394 | 0.483491602 | 0.691473278 | -0.207981675 |

| | | | | | | | | |
|---|---|---|---|---|---|---|---|---|
| chr6 | 31183835 | 31184332 | 8 | 498 | -77.037022 | 0.434236671 | 0.712218678 | -0.277982007 |
| chr6 | 33074380 | 33075732 | 9 | 1353 | 76.84287906 | 0.325637126 | 0.094302941 | 0.231334185 |
| chr6 | 32694246 | 32695986 | 11 | 1741 | 75.00128611 | 0.708990411 | 0.468588051 | 0.240402359 |
| chr6 | 30098965 | 30099981 | 11 | 1017 | -74.7155953 | 0.295544794 | 0.436534629 | -0.140989835 |
| chr6 | 31360626 | 31361977 | 14 | 1352 | 73.49642306 | 0.910506312 | 0.79356845 | 0.116937862 |
| chr6 | 30186993 | 30187774 | 12 | 782 | -72.8472287 | 0.862285347 | 0.972321412 | -0.110036066 |
| chr6 | 31134963 | 31135573 | 11 | 611 | 72.01447718 | 0.763752121 | 0.599935347 | 0.163816774 |
| chr6 | 33097162 | 33098191 | 10 | 1030 | -71.9862418 | 0.33951913 | 0.98066853 | -0.6411494 |
| chr6 | 31359105 | 31359599 | 9 | 495 | 69.06025311 | 0.831645181 | 0.65918805 | 0.172457131 |
| chr6 | 32417193 | 32418725 | 11 | 1533 | -68.6472904 | 0.350978769 | 0.518899593 | -0.167920824 |
| chr6 | 32566799 | 32567547 | 14 | 749 | 67.32311128 | 0.801320076 | 0.647549502 | 0.153770573 |
| chr6 | 33018831 | 33019900 | 11 | 1070 | 63.37111114 | 0.429339357 | 0.318448578 | 0.110890779 |
| chr6 | 31171379 | 31172278 | 13 | 900 | -63.0414094 | 0.723007504 | 0.824586107 | -0.101578603 |
| chr6 | 29958517 | 29959346 | 5 | 830 | -62.8948548 | 0.347234445 | 0.657723969 | -0.310489524 |
| chr6 | 30358555 | 30359193 | 11 | 639 | 61.54463157 | 0.442902519 | 0.332731515 | 0.110171004 |
| chr6 | 31440560 | 31441187 | 15 | 628 | 60.41212278 | 0.601543562 | 0.471649453 | 0.129894109 |
| chr6 | 29960419 | 29961060 | 9 | 642 | -60.0334242 | 0.598602643 | 0.769531839 | -0.170929196 |
| chr6 | 30002552 | 30003896 | 11 | 1345 | 59.90735759 | 0.349911509 | 0.249039891 | 0.100871618 |
| chr6 | 32837365 | 32837793 | 8 | 429 | 59.81729591 | 0.45550256 | 0.278839329 | 0.176663231 |
| chr6 | 33082454 | 33083529 | 12 | 1076 | -59.6557447 | 0.671241049 | 0.804082829 | -0.132841779 |
| chr6 | 31578621 | 31579292 | 12 | 672 | 58.36268744 | 0.5316876 | 0.417037887 | 0.114649713 |
| chr6 | 33001624 | 33002478 | 12 | 855 | 58.28829448 | 0.683623674 | 0.570484468 | 0.113139206 |
| chr6 | 29845704 | 29847183 | 11 | 1480 | 57.51801982 | 0.459444647 | 0.309028254 | 0.150416392 |
| chr6 | 32359950 | 32361341 | 8 | 1392 | 57.32723455 | 0.46924776 | 0.284836795 | 0.184410965 |
| chr6 | 32748015 | 32749374 | 11 | 1360 | 56.08960491 | 0.257024437 | 0.152350409 | 0.104674028 |
| chr6 | 30736750 | 30737032 | 9 | 283 | -55.0016072 | 0.153838999 | 0.342988904 | -0.189149904 |
| chr6 | 32413523 | 32414111 | 13 | 589 | 54.82425259 | 0.208921596 | 0.088753694 | 0.120167902 |
| chr6 | 29793079 | 29793511 | 10 | 433 | 54.07969666 | 0.593841156 | 0.299983209 | 0.293857947 |
| chr6 | 32846087 | 32846545 | 8 | 459 | 52.02973207 | 0.725676531 | 0.575225633 | 0.150450899 |
| chr6 | 29756318 | 29757543 | 11 | 1226 | 50.87994343 | 0.706643376 | 0.601960349 | 0.104683028 |
| chr6 | 29733559 | 29734797 | 8 | 1239 | -48.4226663 | 0.656073919 | 0.802273483 | -0.146199563 |
| chr6 | 32569893 | 32571113 | 8 | 1221 | -48.342276 | 0.657187331 | 0.785785489 | -0.128598158 |

| | | | | | | | | |
|---|---|---|---|---|---|---|---|---|
| chr6 | 30320505 | 30321421 | 10 | 917 | -48.2144284 | 0.698222055 | 0.81196631 | -0.113744255 |
| chr6 | 32578668 | 32579455 | 12 | 788 | 45.87702532 | 0.933207733 | 0.65137405 | 0.281833683 |
| chr6 | 31175520 | 31176139 | 11 | 620 | 45.73183517 | 0.90807978 | 0.802055876 | 0.106023905 |
| chr6 | 31160086 | 31160627 | 9 | 542 | -45.1502315 | 0.555026446 | 0.684723586 | -0.129697139 |
| chr6 | 30804516 | 30805408 | 6 | 893 | -43.9924953 | 0.581769501 | 0.781721331 | -0.19995183 |
| chr6 | 31388065 | 31389214 | 9 | 1150 | -42.4780527 | 0.40631881 | 0.660556563 | -0.254237753 |
| chr6 | 32621968 | 32623532 | 9 | 1565 | -42.3960516 | 0.331817583 | 0.557814029 | -0.225996446 |
| chr6 | 30781889 | 30782146 | 8 | 258 | 41.81995546 | 0.286220263 | 0.140971172 | 0.145249091 |
| chr6 | 33069719 | 33070255 | 9 | 537 | 41.48070622 | 0.918237287 | 0.800772792 | 0.117464495 |
| chr6 | 32851118 | 32852030 | 7 | 913 | -41.3578371 | 0.614043755 | 0.759785094 | -0.145741339 |
| chr6 | 32924606 | 32925903 | 9 | 1298 | -41.2789903 | 0.634941645 | 0.749960262 | -0.115018618 |
| chr6 | 31126493 | 31126880 | 9 | 388 | -40.5871756 | 0.340960426 | 0.450162154 | -0.109201728 |
| chr6 | 29682419 | 29682742 | 10 | 324 | -40.2932798 | 0.532524391 | 0.661616203 | -0.129091813 |
| chr6 | 31128732 | 31129301 | 7 | 570 | -40.2888145 | 0.174413535 | 0.315739838 | -0.141326303 |
| chr6 | 33051956 | 33052575 | 8 | 620 | -40.0612748 | 0.412726904 | 0.580517166 | -0.167790262 |
| chr6 | 30356126 | 30356340 | 6 | 215 | 39.2440906 | 0.460044021 | 0.31825397 | 0.14179005 |
| chr6 | 29730626 | 29731089 | 9 | 464 | 39.1630445 | 0.76381997 | 0.624502975 | 0.139316995 |
| chr6 | 32568374 | 32568824 | 6 | 451 | -38.1961603 | 0.582178465 | 0.727210183 | -0.145031718 |
| chr6 | 32545913 | 32546524 | 7 | 612 | -37.791585 | 0.365627584 | 0.532851038 | -0.167223454 |
| chr6 | 32261259 | 32261698 | 8 | 440 | 37.71134419 | 0.532905641 | 0.412307204 | 0.120598436 |
| chr6 | 28962189 | 28962661 | 9 | 473 | 36.79426237 | 0.433592695 | 0.325820253 | 0.107772442 |
| chr6 | 31021025 | 31021108 | 10 | 84 | -36.3547136 | 0.397738568 | 0.514009239 | -0.116270671 |
| chr6 | 28958730 | 28958996 | 8 | 267 | 35.27403535 | 0.443244438 | 0.323304033 | 0.119940404 |
| chr6 | 32027971 | 32028258 | 9 | 288 | 35.17603184 | 0.413987387 | 0.294821078 | 0.119166309 |
| chr6 | 32390385 | 32391056 | 5 | 672 | -35.172679 | 0.233048879 | 0.442913747 | -0.209864867 |
| chr6 | 32471318 | 32471954 | 8 | 637 | -34.8278199 | 0.662966562 | 0.775189548 | -0.112222985 |
| chr6 | 31458220 | 31459412 | 7 | 1193 | -34.6541987 | 0.235634028 | 0.421615662 | -0.185981634 |
| chr6 | 29634996 | 29635486 | 7 | 491 | 34.63033415 | 0.481242807 | 0.350232817 | 0.13100999 |
| chr6 | 33159608 | 33159810 | 6 | 203 | 34.15112373 | 0.415209162 | 0.274221686 | 0.140987476 |
| chr6 | 30792650 | 30792755 | 9 | 106 | -33.6435716 | 0.818058636 | 0.919958424 | -0.101899787 |
| chr6 | 32720456 | 32721248 | 6 | 793 | 33.63610016 | 0.396183486 | 0.251808767 | 0.144374719 |
| chr6 | 30897463 | 30897689 | 8 | 227 | 33.01332089 | 0.847765094 | 0.737263331 | 0.110501763 |

| | | | | | | | | |
|---|---|---|---|---|---|---|---|---|
| chr6 | 31201857 | 31202353 | 5 | 497 | -32.1758323 | 0.513265733 | 0.660753211 | -0.147487478 |
| chr6 | 32536976 | 32537593 | 7 | 618 | -30.6116319 | 0.585999172 | 0.77951717 | -0.193517998 |
| chr6 | 31141990 | 31142187 | 7 | 198 | -29.9808869 | 0.698629217 | 0.801718357 | -0.103089139 |
| chr6 | 32719125 | 32719789 | 6 | 665 | 29.60650157 | 0.411442301 | 0.274916962 | 0.136525339 |
| chr6 | 31556619 | 31556748 | 5 | 130 | -29.1536559 | 0.171624347 | 0.332976159 | -0.161351811 |
| chr6 | 32693389 | 32693515 | 3 | 127 | 29.05072735 | 0.569743794 | 0.265661451 | 0.304082343 |
| chr6 | 32741693 | 32742432 | 6 | 740 | 29.0450097 | 0.328939094 | 0.203656213 | 0.125282881 |
| chr6 | 29612525 | 29613261 | 7 | 737 | 28.65052131 | 0.485811718 | 0.329570373 | 0.156241345 |
| chr6 | 29625867 | 29626704 | 5 | 838 | 28.30574815 | 0.499008003 | 0.334341324 | 0.164666679 |
| chr6 | 29981380 | 29981719 | 7 | 340 | 27.6748833 | 0.722203514 | 0.57345365 | 0.148749864 |
| chr6 | 32469092 | 32470024 | 6 | 933 | 27.28727006 | 0.856146292 | 0.632055921 | 0.224090371 |
| chr6 | 30393065 | 30393731 | 5 | 667 | -27.2349303 | 0.322830738 | 0.438911965 | -0.116081227 |
| chr6 | 30886456 | 30886774 | 4 | 319 | 26.78454321 | 0.595374945 | 0.435961337 | 0.159413608 |
| chr6 | 31354443 | 31354736 | 7 | 294 | 26.159898 | 0.393398028 | 0.173403791 | 0.219994237 |
| chr6 | 31430051 | 31430765 | 6 | 715 | 25.63288391 | 0.328049039 | 0.210621966 | 0.117427073 |
| chr6 | 29927297 | 29927339 | 6 | 43 | 25.00896438 | 0.557886836 | 0.392595212 | 0.165291624 |
| chr6 | 30104839 | 30105998 | 5 | 1160 | -24.6384274 | 0.393195101 | 0.510534072 | -0.117338971 |
| chr6 | 31664319 | 31664471 | 5 | 153 | -24.507382 | 0.45716527 | 0.560084713 | -0.102919443 |
| chr6 | 30858820 | 30859129 | 5 | 310 | 23.80352118 | 0.472786476 | 0.361402832 | 0.111383644 |
| chr6 | 30793441 | 30793795 | 5 | 355 | 23.62831899 | 0.382616179 | 0.225878375 | 0.156737803 |
| chr6 | 29944116 | 29944308 | 6 | 193 | -23.4002679 | 0.479542339 | 0.619952462 | -0.140410123 |
| chr6 | 31483502 | 31484149 | 5 | 648 | 23.37490789 | 0.77174244 | 0.489952985 | 0.281789455 |
| chr6 | 32580169 | 32580776 | 6 | 608 | 22.35398813 | 0.943452289 | 0.59831769 | 0.345134599 |
| chr6 | 32663478 | 32663807 | 4 | 330 | -22.3120384 | 0.515751077 | 0.688478666 | -0.172727589 |
| chr6 | 31276436 | 31276993 | 4 | 558 | 21.88182005 | 0.553897451 | 0.445827166 | 0.108070285 |
| chr6 | 29799445 | 29799617 | 5 | 173 | -21.4273707 | 0.464729269 | 0.578315484 | -0.113586214 |
| chr6 | 30806909 | 30807103 | 5 | 195 | -20.843034 | 0.820574837 | 0.930630748 | -0.11005591 |
| chr6 | 32386030 | 32386368 | 5 | 339 | 20.54744387 | 0.639283041 | 0.484369739 | 0.154913301 |
| chr6 | 33233833 | 33234617 | 5 | 785 | 20.38373728 | 0.458354919 | 0.35330735 | 0.105047569 |
| chr6 | 29720358 | 29720508 | 3 | 151 | -20.348127 | 0.464002244 | 0.766190486 | -0.302188242 |
| chr6 | 30029837 | 30030323 | 4 | 487 | 19.85184045 | 0.826767006 | 0.724674397 | 0.10209261 |
| chr6 | 33100404 | 33100505 | 3 | 102 | -19.8201156 | 0.241831579 | 0.891165688 | -0.649334109 |

| | | | | | | | | |
|---|---|---|---|---|---|---|---|---|
| chr6 | 30887405 | 30887851 | 3 | 447 | 17.64754233 | 0.713873705 | 0.569845617 | 0.144028088 |
| chr6 | 33058539 | 33058775 | 4 | 237 | -17.4291771 | 0.457032311 | 0.966089847 | -0.509057536 |
| chr6 | 33008465 | 33008795 | 3 | 331 | 17.19809934 | 0.356927525 | 0.237492934 | 0.119434591 |
| chr6 | 29753780 | 29753939 | 3 | 160 | -16.8662142 | 0.300010503 | 0.425324465 | -0.125313962 |
| chr6 | 29941060 | 29941199 | 4 | 140 | -16.810257 | 0.397358218 | 0.51956924 | -0.122211021 |
| chr6 | 30799343 | 30799637 | 4 | 295 | -16.7036026 | 0.690178401 | 0.796508089 | -0.106329688 |
| chr6 | 30112865 | 30113331 | 4 | 467 | -16.3809398 | 0.264793004 | 0.400075746 | -0.135282741 |
| chr6 | 29530926 | 29531639 | 4 | 714 | 16.02521482 | 0.276601805 | 0.081729281 | 0.194872524 |
| chr6 | 30736448 | 30736480 | 4 | 33 | 16.00370432 | 0.504137152 | 0.398735219 | 0.105401934 |
| chr6 | 32763350 | 32763550 | 4 | 201 | 15.94407524 | 0.19520521 | 0.063673709 | 0.131531501 |
| chr6 | 29441981 | 29442290 | 3 | 310 | 15.88709338 | 0.384589667 | 0.278410384 | 0.106179283 |
| chr6 | 29975878 | 29975911 | 3 | 34 | 15.85735395 | 0.640100756 | 0.447922901 | 0.192177855 |
| chr6 | 32271885 | 32272516 | 4 | 632 | 15.60859711 | 0.609644709 | 0.505679411 | 0.103965298 |
| chr6 | 33030296 | 33030361 | 3 | 66 | -15.2127155 | 0.041269154 | 0.190702284 | -0.14943313 |
| chr6 | 29016306 | 29016457 | 4 | 152 | -15.1186513 | 0.452252172 | 0.591049777 | -0.138797605 |
| chr6 | 31131375 | 31131465 | 4 | 91 | 14.43964552 | 0.59157452 | 0.483172529 | 0.108401991 |
| chr6 | 32482295 | 32482719 | 3 | 425 | -12.7222333 | 0.572894843 | 0.976884588 | -0.403989745 |
| chr6 | 29754814 | 29754927 | 3 | 114 | 12.49430183 | 0.81618596 | 0.708016385 | 0.108169576 |
| chr6 | 29890583 | 29890883 | 3 | 301 | -12.4837022 | 0.792020976 | 0.894029776 | -0.1020088 |
| chr6 | 30170867 | 30171118 | 3 | 252 | 12.29848739 | 0.528483277 | 0.42240026 | 0.106083017 |
| chr6 | 30815343 | 30815600 | 3 | 258 | 11.8809758 | 0.784192467 | 0.660893733 | 0.123298733 |
| chr6 | 28971625 | 28971685 | 3 | 61 | 11.8720525 | 0.674696673 | 0.516988175 | 0.157708497 |
| chr6 | 29700471 | 29700968 | 3 | 498 | -11.7898692 | 0.274710582 | 0.519870163 | -0.24515958 |

### Supplementary Table 6. Sequences of sgRNAs and PCR primers

| | Name | DNA sequence (5'-3') |
|---|---|---|
| **sgRNA set1** | TR1 | AGGCTGGGGTAGGCCGTGTC |
| | TR2 | AGACTGAAGTAGGGCCGGAC |
| | TR3 | GTTGGCTGTAATCCTACGCC |
| | TR4 | CATGTTCCACGCCCTTGCAG |
| | TR5 | ACTGTTCCCAGACGGAAAAC |
| | TR6 | CTGGTTTTACACATCGCTGG |
| | DD1 | TGGATAGAGGAGGCTTATAC |
| | DD2 | TCGTTGCTCAGGTGGTGGTA |
| | DD3 | TGGATGGGTTCTCGCAAAAT |
| | DD4 | AGCCACAGTCGACAATGGCC |
| | PS1 | AAGATGCATTGAGCTCTGCG |
| | PS2 | GGCAGAGCCCCGTGCATCCT |
| | PS3 | GGTAGCCTGAGACTTCGTAA |
| | PS4 | TTCCAACAGGAGTCTTACCT |
| | ZBT1 | GCAGGAACCCTTCGAGCCTA |
| | ZBT2 | GAGCAGGCCAGGCCGTGCAT |
| | ZBT3 | ATGATGTGCCGGTCACGCTT |
| | ZBT4 | AGGCATGGACTCGGAACCGA |
| **sgRNA set2** | ZBE1 | TCTTTCTGTCACTCCTAAGG |
| | ZBE2 | TGAGTTGTTAGTTTGCATCC |
| | ZBE3 | AAACTGAGGATATCCGAAAG |
| | ZBE4 | ATAAATCTGTACTCCCCATC |
| | ZBE5 | ACATGGTGGACGTACTCGCA |
| | VA1 | TCTTTTCACCGGGTTTCGTA |
| | VA2 | CAGGCAGCCTCAACATATCG |
| | VA3 | CCCCTCTCACCAGCGCACGA |
| | VA4 | TGTTGCAGGCACCGGATGCG |
| | VA5 | GAAGGAGGCGTGAGTATGAT |
| | VP1 | GTGGATGGTAAGCGATGGCA |
| | VP2 | CTGGAGGATACAATCCCGAA |

| | | |
|---|---|---|
| | VP3 | CTGCTCCATTCGCTGTAGGG |
| | VP4 | GAAATCGCCAGGCAGTTCGG |
| | VP5 | ATGTCAGAGGCGTGCTCGAT |
| | VP6 | GTCTTACTACCGCTCTTACC |
| **PCR primers for the amplication of the sgRNA targeted region** | ZBED9-F | CTACCGTCGCTTGGCTAGAT |
| | ZBED9-R | GAGGGCACTCCACAAATTGC |
| | ZBTB9-F | TGTCATTGAAGCCGATGCCT |
| | ZBTB9-R | CTTTCCTGCAGGGTTGGCTA |
| | VARS2-F1 | GCAGGGGAGAGCAAGGTTAG |
| | VARS2-R1 | ACCAAGGCACAGAACACTCA |
| | VARS2-F2 | CTCTCGCCTCTTTTCGACCA |
| | VARS2-R2 | GAAGTCAGAGCTGTGTCGGG |
| | VPS52-F1 | TCACTGGATCGTCTCACTCCT |
| | VPS52-R1 | CAGCCCATCCACCTGCTATG |
| | VPS52-F2 | CAGAGCCCAGGTTCTTGGAG |
| | VPS52-R2 | GGAGTCGAAAGTCCTCCCAC |
| **qPCR primers** | HFE-F | TGGGACGTGGCTAGTCATAA |
| | HFE-R | CATCCCAAATGAAGGCACCATTA |
| | MUC21-F | TGGTTAAAATTGGGCTCTGG |
| | MUC22-R | GCAGCCGACTCTTCTTGTTC |
| | TCF19-F | ATGGAGCCCAGGAAGAAACT |
| | TCF19-R | TAACCCAGGTCTTGCCATTC |
| | MICB-F | TTCCACCCCTCTTCTTGCTA |
| | MICB-R | CCCACAACCCTGTATTGGAC |
| | RNF8-F | TCTCTCCAACTGCCTGCTTG |
| | RNF9-R | GGCCCCACACCTATGAACAA |
| **HLA-C Sanger sequencing** | C565-out-F | CGCAGTCCCGGTTCTAAAGT |
| | C565-out-R | TCTACGGGAGATGGGGAAGG |
| | C565-in-F | CGCGGATCCTCTTCACATCCGTGTCCCG |
| | C565-in-R | CCGGAATTCGGTAAAGGTGACTGGGGCTC |

| | | |
|---|---|---|
| | C1625-out-F | AGGAGGGGAGGAAAATGGGA |
| | C1625-out-R | GAGGTGGGGCACACTTCTAC |
| | C1625-in-F | CGCGGATCCGTCAGGCTGCTGACCTTTCT |
| | C1625-in-R | CCGGAATTCTACCCCATCTCAGGGTGAGG |
| **ASE pyro sequencing** | DPA1Eout-F | GCACAGTCTCCGTTGTCTCA |
| | DPA1Eout-R | CTGTAGCCCAACTGGAAGGG |
| | DPA1Ein-F | GCAGAGGACACCAGGTCTTT |
| | DPA1Ein-R | /5Biosg/CAGAGTGAGGCTGGTCAGTG |
| | DPA1-C-out-F | TTTATGATGAGGACGGTGCCC |
| | DPA1-C-out-R | TTGATCCAGCGTTCCAACCA |
| | DPA1-C-in-F | GAGGGCACAAAGGTCAGGTAAT |
| | DPA1-C-in-R | /5Biosg/ACACCCTCATCTGCCACATTG |
| | DPA1seq | CTCAGCGACACCCTCAGTG |
| **ASM Sanger sequencing** | DPA1Mout-F | TTGTTGTTGGGTAAAGAGGA |
| | DPA1Mout-R | AACCCCACCTCCTTCAAAA |
| | DPA1Min-F | CGCGGATCCGATTTTGTTTTTAAGAAGGG |
| | DPA1Min-R | CCGGAATTCAACCACCAATTCTAATTTCA |

/5Biosg/ = 5’ biotin added
